# Seipin and perilipin 3 promote distinct early steps of nascent lipid droplet biogenesis

**DOI:** 10.64898/2026.10.21.760741

**Authors:** Raunaq Deo, Apoorva Bhapkar, Emma Givan, Chao Li, Ehud Haimov, Derk Binns, Philipp E Scherer, Michael M. Kozlov, Alexandre Toulmay, William A. Prinz

## Abstract

Lipid droplets (LDs) are lipid storage organelles with critical roles in metabolism. Cells form LDs in response to various metabolic stresses^1^. LD biogenesis occurs in the endoplasmic reticulum (ER) membrane and begins when triglyceride (TG) in the membrane phase separates^2–4^. After nucleation, nascent LDs grow by absorbing TG from the membrane^5,6^. How cells determine when TG in the ER membrane is added to existing LDs or is used to form new LDs or is not known. Here, we developed the first LD formation assay that allows the synthesis of TG in membranes, mimicking what occurs in cells. LDs arise stochastically after TG production begins. LD nucleation is affected by TG production rate and membrane phospholipid composition. Membrane-embedded LDs suppress nascent LD formation by sequestering TG and destabilizing nascent LDs by Ostwald ripening. We show that cells overcome this suppression in two ways. The ER membrane protein seipin accelerates LD nucleation where it localizes in a membrane, probably by increasing the kinetics of TG phase separation. The LD coat perilipin 3 inhibits nascent droplet dissipation by ripening. Together, our findings reveal a minimal LD formation machinery, and the mechanisms cells use to control whether TG in the ER drives new LD formation.

## Main

Lipid droplets (LDs) are ubiquitous storage organelles that function as critical protective lipid buffers, blunting lipotoxicity by sequestering excess and damaged lipids^1,7^. Accordingly, cells tightly regulate the formation of LDs in response metabolic stresses. How cells determine whether to add excess lipid to existing LDs or use it to form new LDs is not well understood.

LDs have a core of neutral lipids, mostly triglyceride (TG) and steryl esters, surrounded by a monolayer of phospholipids that contains proteins. They form in the endoplasmic reticulum (ER), where neutral lipid synthesis occurs^1^. After LDs form, most remain embedded in the ER and have a surface phospholipid monolayer that is continuous with the cytosolic leaflet of the ER membrane^8–12^. This architecture enables the movement of TG from the ER membrane into existing LDs^13^.

The first step of LD formation is thought to be driven by TG phase separation in phospholipid bilayers^2,4,6^. TG is poorly soluble in membranes. The solubility limit is called the critical concentration; above this limit, TG phase separates (demixes) into an oil phase between the two leaflets of a membrane^4^. Studies using artificial LDs^14–18^ or molecular dynamics (MD) simulations^5,19,20^ indicate demixing is spontaneous and depends on the TG concentration and phospholipids in the membrane, which provide the energy barrier to TG demixing^4,6,19,21–23^. However, in the experimental systems used for these studies, TG is incorporated into membranes from the start and is not enzymatically synthesized in the membrane, as it is in cells. When TG is produced in membranes, the determinants of nascent LD formation and growth remain unknown.

Several proteins have been implicated in nascent LD biogenesis. The best characterized is seipin, which is required for proper LD formation and lipid storage^24–27^. Seipin forms an oligomeric ring^28–32^ and is localized to ER-LD junctions^33,34^. In silico^35–37^ and biochemical experiments^26,31,38^ suggest TG molecules clustering within seipin is a prelude to TG phase separation, though the mechanism of nucleation remains unresolved. Mammalian seipin is regulated by two proteins that associate with it: Lipid Droplet Assembly Factor 1 (LDAF1) and adipogenin (Adig). LDAF1 has been suggested to facilitate LD biogenesis together with seipin by increasing TG binding or altering TG diffusion within the seipin oligomer^18,39,40^. The combination of seipin and LDAF1 has been termed the Lipid Droplet Assembly Complex (LDAC). On the other hand, Adig is thought to inhibit the ability of seipin to promote LD formation^41^. Another putative LD biogenesis protein is perilipin 3 (PLIN3), the first protein to become enriched on nascent LDs after LDAC^39^. PLIN3 (and other perilipins) coats the surface of mature LDs, regulating the access of enzymes to the stored lipids^42^, but its contribution to LD formation is unknown. It should be noted that LDs still form in cells lacking seipin, LDAF1, Adig or PLIN3. It has been suggested that LDs formed without these proteins do not function optimally^43,44^.

Here, we reconstitute LD formation in model membranes by inducing TG synthesis, as occurs in cells. Our analysis reveals how nascent LD nucleation depends on the kinetics of TG phase separation. We show how increasing TG synthesis or altering the phospholipid composition of the ER membrane can promote nascent LD formation *in vitro* and in cells. We find pre-existing membrane-embedded LDs are indeed sinks for TG and suppress nascent LD formation. This suppression is overcome by seipin, which accelerates TG nucleation, and PLIN3, which stabilizes TG within nascent LDs once they form.

### TG synthesis in a bilayer is sufficient to drive nascent LD formation

To reconstitute LD formation, we first generated supported lipid bilayers (SLBs) on glass coverslips in a flow cell by fusing liposomes^45,46^ composed of dioleoyl phosphatidylcholine (DOPC) and 10 mol% dioleoyl glycerol (DG) (Fig. 1A, Fig. S1A). A soluble bifunctional wax synthase/diacylglycerol acyltransferase (sDGAT) from *Acinetobacter baumannii* was then used to synthesize TG from the DG in the SLBs (Fig. S1B). This enzyme only catalyzes the formation of TG and not wax esters when provided with DG and a fatty acyl-CoA^47^(Fig. S1C). We recruited sDGAT to SLBs by engineering a 6X His-tag at its N-terminus and using SLBs with 2.5 mol% of the nickel-containing lipid DOGS-3-[(N-(5-amino-1-carboxypentyl) iminodiacetic acid) succinyl] (NiNTA). TG synthesis was initiated by introducing oleoyl-CoA to the flow cell.

**Figure 1:**
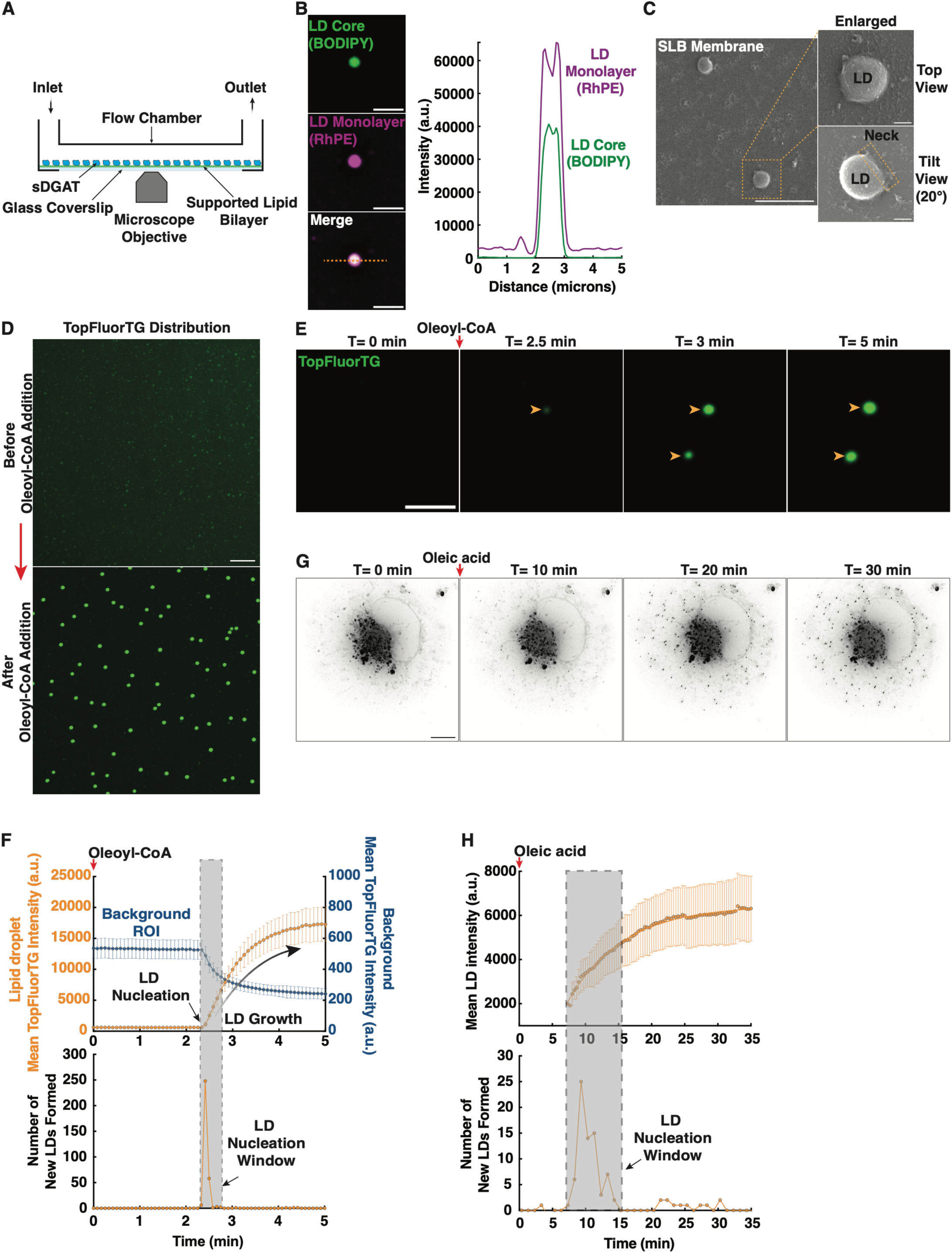
TG synthesis is sufficient to drive LD formation. A. Cartoon of flow cells used to quantify LD formation in SLBs in real time. B. SLBs with DOPC:DG:DOGS-NiNTA:RHPE (87.3:10:2.5:0.2 mol%) were incubated with 1µM sDGAT and 18µM oleoyl-CoA at 30°C for one hour and stained with BODIPY (left). Bar = 2 μm. Fluorescence signal along dotted line (right). C. SEM images of droplets formed on SLBs (top view and tilted view, 20° tilt, of one droplet enlarged). Bar = 2 μm, enlarged image = 200nm. D. SLBs membranes with DOPC:DG:DOGS-NiNTA:TopFluorTG (87.4:10:2.5:0.1 mol%) before (top) and after 5 min incubation with 1µM sDGAT and 18µM oleoyl-CoA (bottom). Bar = 5 μm. E. SLBs membranes as in panel D were imaged every 5 seconds after oleoyl-CoA addition (T=0). Images from Movie 1; arrowheads indicate nascent LDs. Bar = 2 μm. F. Top: mean intensity of ∼300 LDs over time from experiment as in Fig. 1E. Background calculated from region of SLB where LDs did not form. Mean +/− std. dev. of images taken every 5 seconds. Bottom: number of LDs formed over time in the reaction. Grey bar indicates nucleation window. G. Images of a Cos-7 cell stained with BODIPY, before and after addition of 400uM oleic acid. Images acquired every 20s. Bar = 10 µm. H. Quantification of experiment in 1G. Top: mean intensity of the 87 LDs that form after oleic acid addition. Mean +/− std. dev. of images taken every 20 seconds. Bottom: number of LDs formed over time in the reaction. Grey bar indicates nucleation window.

We included trace amounts of rhodamine-labeled phosphatidylethanolamine (RHPE) in the SLBs to label the membrane (Fig. S1A). To observe LDs after the reaction, we added dipyrrometheneboron difluoride (BODIPY) 493/503, a lipophilic dye that labels LD cores^48^. One hour after oleoyl-CoA addition, we observed BODIPY-labeled puncta encircled by RHPE, consistent with LDs covered by a phospholipid monolayer (Fig. 1B). Fluorescence recovery after photobleaching (FRAP) confirmed that RHPE diffused freely between the bilayer and LDs. This indicates LDs remain membrane-embedded after formation and do not detach (Fig. S1D). We further characterized the ultrastructure of these LDs using scanning electron microscopy (Fig. 1C), which revealed droplets attached to the SLBs (Fig. 1C, bottom right). LDs had a mean diameter of 0.52 +/− 0.08 µm (n=17). These experiments show we established an *in vitro* assay of LD formation.

To visualize and quantify the kinetics of nascent LD formation in real time, we used SLBs containing trace amounts (0.1 mol%) of TopFluorTG, a fluorescent TG analog that accumulates in LDs ^15,16,18^. We confirmed that before TG production was initiated TopFluorTG was uniformly distributed and diffused freely within the bilayer (Fig. S1E). We then used time-lapse fluorescence microscopy to capture LD formation events in real time after oleoyl-CoA was added to the flow cell (Movie 1, Fig. 1D). Note that it takes about 1.3 minutes for the oleoyl-CoA to reach the SLB in the flow cell (not shown). Two and half minutes after oleoyl-CoA was added to the flow cell, we observed bright TopFluorTG puncta, LDs, on the SLBs (Fig. 1E). This indicates that as TG is synthesized in an SLB it undergoes phase separation, likely after exceeding the critical concentration of TG in the membrane, nucleating nascent LD formation (Fig.1E; Movie 1). As LDs form, TopFluorTG is rapidly depleted from the surrounding bilayer and accumulates in nucleated droplets (Movie 1; Fig. 1F, top). Quantifying mean TopFluorTG fluorescence for ∼300 LDs over time revealed that ∼83% of the nucleation events occurred within a short (∼25 second) nucleation window (Fig. 1F bottom; Movie 1). After nucleation, the fluorescence signal of newly formed LDs continued to increase, indicating these LDs continue to grow by acquiring additional TG from the membrane (Fig. 1F). Note that we found a linear relationship between LD fluorescence intensity and LD size as assessed by fluorescence microscopy (Fig. S1F), indicating that measuring mean intensity is an effective method of estimating relative LD size. The formation of LDs is not affected by the location of sDGAT; we found it does not become enriched on nascent LDs (Fig. S1G, H).

Although LDs appear to form stochastically within the SLBs, we asked whether they preferentially nucleate at regions of the SLB where unfused liposomes remain on the surface^45,49^ (Fig. S1A, S1I). To test this, we calculated Manders’ coefficient to quantify the co-occurrence of unfused liposomes and LD formation sites. The coefficient was 0.040 ± 0.007 and 0.037 ± 0.007 after the image of LDs was rotated 90°, indicating no correlation between the location of unfused liposomes on the SLBs and LD formation sites (Fig. S1J).

We found LDs formed on SLBs do not diffuse laterally and do not detach over time. We considered whether these properties might reflect an influence of the underlying glass substrate on LD behavior. To test this, we repeated the LD formation assays using free-standing giant unilamellar vesicles (GUVs) instead of SLBs. As observed on SLBs, multiple LDs nucleated on the GUV membranes (Fig. S1K, Movie 2). We verified that the TopFluorTG puncta on the GUVs are LDs by showing RHPE surrounding these LDs (Fig. S1L, bottom), similar to LDs formed on SLBs (Fig. 1B, right). On GUVs, some smaller LDs initially exhibited lateral mobility along the membrane but became progressively less mobile as they grew larger (Movie 2). Notably, the LDs formed on GUVs, like those on SLBs, did not detach from the membrane. These results suggest that while the glass support under SLBs may restricts LD diffusion, it does not substantially influence LD formation. Collectively, these experiments show that we have established a new *in vitro* system to quantitatively assess the early steps of LD formation and that TG synthesis in a membrane is sufficient to drive LD formation.

### Membrane-embedded droplets suppress nascent LD formation

Our findings suggest that once LDs form in SLBs they are sinks for TG in the membrane and consequently suppress subsequent nucleation events. To determine whether this occurs in cells too, we followed LD formation and growth in COS-7 cells after oleic acid was added to them, which is known to trigger nascent LD biogenesis^25^. Fig. 1G shows images of a representative cell over time (Movie 3). We tracked the nucleation and growth of all 84 LDs that formed in this cell within 35 minutes of oleic acid addition (Fig. 1H). We observed that ∼87% of these LDs appear within a 7 min nucleation window (Fig H1, bottom) and then continue to grow (Fig. 1G, Fig. H1, top). Note that no LD coalescence events were observed. Therefore, our model membrane-based LD formation system recapitulates what occurs in cells and suggests that, in both cells and SLBs, LDs that nucleate first suppress subsequent nucleation events, probably by absorbing TG from the membrane and preventing the TG concentration from rising above the critical concentration.

To further investigate how membrane-embedded LDs affect nascent LD formation, we generated LDs in SLBs by two rounds of TG synthesis. In the first, LDs were generated by adding 1.8 µM oleoyl-CoA, resulting in partial DG consumption. We refer to these as pre-existing LDs. We then added 18 µM oleoyl-CoA, triggering a second round of TG synthesis. Interestingly, all the LDs formed in the second round of LD production shrank and eventually disappeared while all pre-existing LDs grew (Movie 4; Fig. 2A, B). This phenomenon is characteristic of Ostwald ripening, which has been observed to drive TG redistribution between droplets^16,26^. We also used FRAP to confirm that TopFluorTG in the LDs is mobile, consistent with the idea that TG can redistribute between LDs as required for ripening^26^(Fig. 2C).

**Figure 2.**
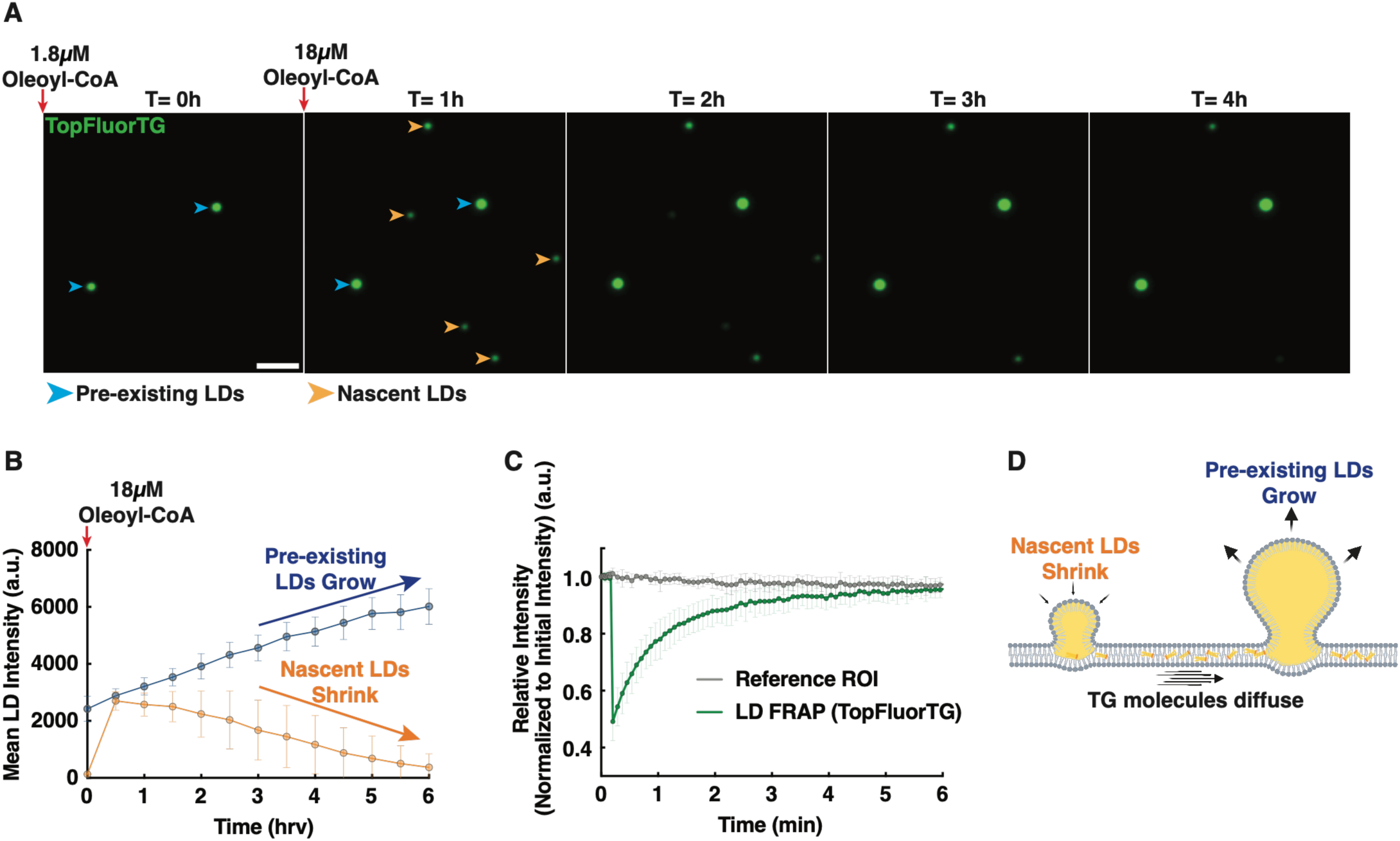
Pre-existing LDs suppress new LD formation by Ostwald ripening. A. Pre-existing LDs were generated in SLBs with 10 mol% DG by adding 1.8 <u>µ</u>M oleoyl-CoA for 1 hour. This results in LD formation but does not deplete all DG in the SLB. A second round of LD production was initiated by adding 18 µM oleoyl-CoA at time = 0 and images were taken every 10 minutes for 6 hr. Pre-existing LDs indicated with blue arrowheads and nascent LDs that form in the second round of LD production with orange arrowheads. Bar = 2μm. B. Quantification of fluorescence intensity of LDs in experiments as in panel A; mean and std. dev. of 55 pre-existing LDs and 162 nascent LDs. Note that all LDs formed in the second round of LD formation shrink by Ostwald ripening. C. FRAP of TopFluorTG in LDs formed on SLBs with 10 mol% DG. Mean +/− std. dev. of 3 experiments. D. Schematic representation of Ostwald ripening affecting membrane-embedded LDs. Schematic created using BioRender (BioRender.com).

Together, our findings show how membrane-embedded LDs suppress nascent LD formation and suggest the following scenario. The first round of TG synthesis leads to stochastic LD nucleation. After the LDs form, a thermodynamic equilibrium is established between TG molecules in the membrane and the LDs. This equilibrium is transiently disturbed by the second round of TG synthesis, allowing nascent LDs to form even though much of the TG made in the second round is absorbed by the pre-existing LDs (as indicated by their growth). The LDs formed in the second round of TG synthesis are smaller than the pre-existing LDs and undergo Ostwald ripening, where smaller LDs dissolve and shrink while bigger LDs grow. Together, our findings suggest membrane-embedded LDs suppress new LD formation by absorbing TG and destabilizing LDs by Ostwald ripening.

### Velocity of TG production affects the probability of LD nucleation

We sought to understand the relationship between TG production rate and the kinetics of TG phase separation. To start, we used Monte Carlo-like simulations that treat a membrane as a grid of possible LD nucleation sites. The simulations assume that TG is added to the membrane at a constant rate and distributed uniformly throughout the grid. At each time point of the simulation, the probability that TG undergoes phase separation at a nucleation site is assumed to be proportional to the exponential of the energy barrier, which is a function of TG concentration, divided by the thermal energy (Boltzmann factor; see Methods for details). Once LDs form, they absorb free TG molecules at a rate proportional to their size and to the difference in TG chemical potential between the droplet and free environment. Simulations show that this absorption eventually outpaces the TG production rate, thus inhibiting new droplet formation and resulting in a finite number of LDs. Using these simulations, we found that the number of LD nucleation events is proportional to the velocity of TG addition to the membrane (Fig. S2A). This is because the further levels of TG in a membrane increase above the critical concentration, the higher the probability that nucleation will occur at the nucleation sites in the simulated membranes.

We next determined whether the velocity of TG production similarly affects LD formation rates in SLBs. TG production rates were varied by using SLBs containing 10 mol% DG with different concentrations of sDGAT (Fig. 3A). LD formation reactions were allowed to proceed to completion (1 hour). We found that increasing sDGAT concentration produced more LDs (Fig. 3 A, B), matching our Monte Carlo simulations (Fig. S2A). LD nucleation occurred earlier in reactions with high sDGAT (1000 nM); nucleation began ∼2 minutes after oleoyl-CoA addition in reactions with high sDGAT and after ∼22 minutes with low sDGAT (10 nM). (Fig. 3C, S2B). In the reactions with high sDGAT, LDs nucleate within a short ∼25 second window, while LD nucleation continued for about 200 seconds during the reactions with low sDGAT (Fig. 3D). Despite the shorter nucleation window, under high sDGAT conditions about 55 LDs formed per minute per field of view (8281 um^2^), while the nucleation rate was 25-fold slower with low sDGAT. (Fig. 3D). Consequently, 35-fold more LDs formed in the reactions with high sDGAT (Fig. 3D). Once LDs formed, they grew more rapidly in reactions with high sDGAT (Fig. S2C). Note that the LDs formed with high sDGAT were significantly smaller than those formed with low sDGAT (Fig. 3A, S2D). This is probably because TG produced in the high sDGAT reactions was distributed among more LDs. These experiments establish how velocity of TG production influences the kinetics of TG phase separation, and therefore the probability of LD nucleation in our assay.

**Figure 3:**
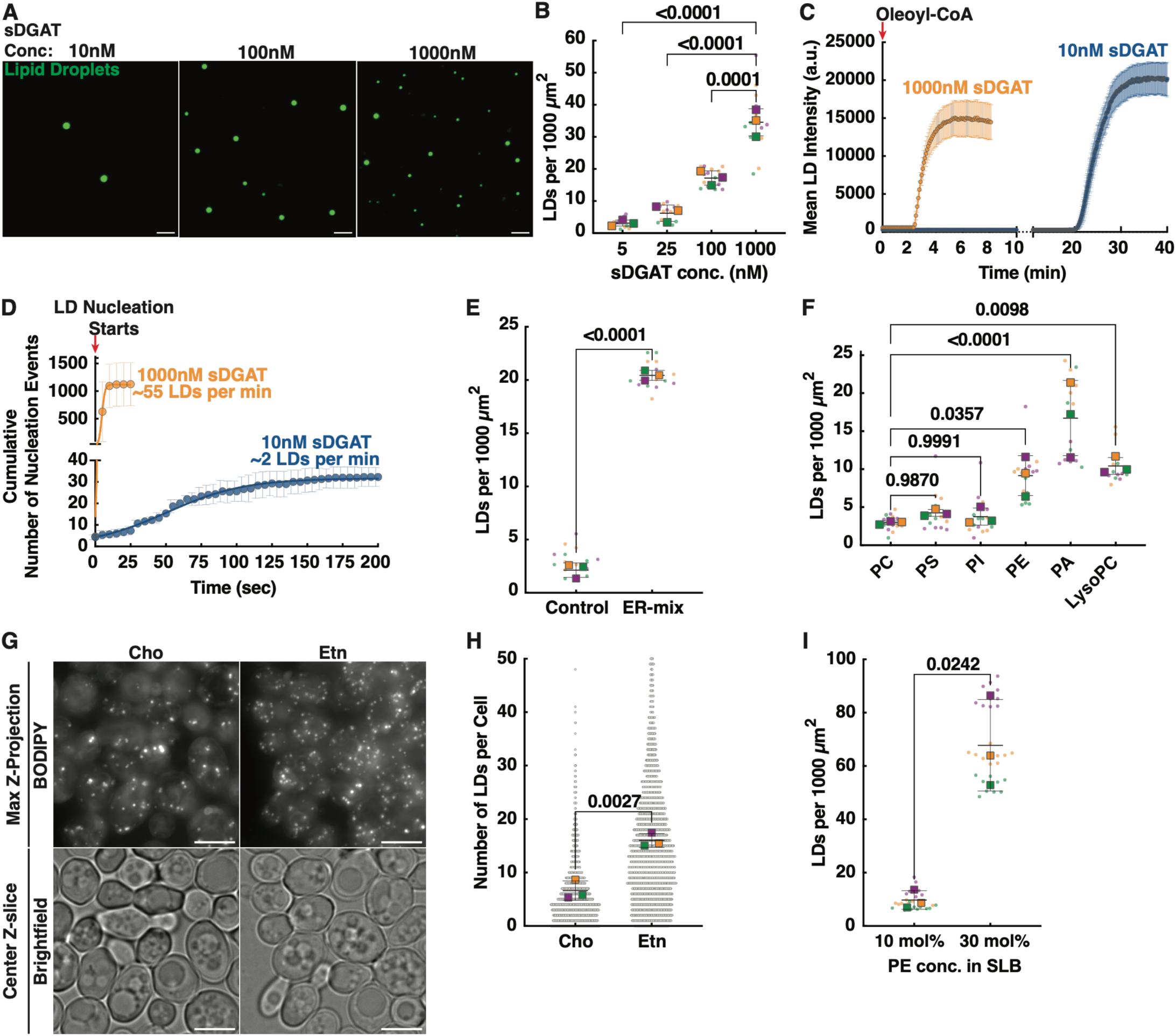
Rate of TG production and membrane phospholipid composition affects LD nucleation. A. LD formation reactions performed on SLBs containing DOPC:DG:DOGS-NiNTA: TopFluorTG (87.4:10:2.5:0.1 mol%) and the indicated concentration of sDGAT. Images taken after 1hr incubation with oleoyl-CoA. Bars = 2μm. B. Density of LDs formed on SLBs in assays as in panel A. Colored squares show the means of independent experiments; dots with the same color are quantifications of single visual fields used to calculate the means. Bars indicate mean and std. dev. of 3 independent experiments; p values calculated from one-way ANOVAs. C. Mean intensity over time of LDs formed on SLBs as in panel A. Oleoyl-CoA was added at time = 0. Mean +/− std. dev. of 10 LDs (10 nm sDGAT) and 295 LDs (1000 nm sDGAT). D. Cumulative number of LDs formed in assays as in panel A. Mean +/− std. dev. from 3 independent experiments. E. Density of LDs formed on SLBs for assays with an ER-like lipid mix or DOPC and 4 mol% DG (control), and 1000 nM DGAT after 1hr incubation (lipid mixes described in Fig. S3A). Colored squares show the means of independent experiments; dots with the same color are quantifications of single visual fields used to calculate the means. Bars are the mean and std. dev. of 3 independent experiments; p value calculated from Welch’s t-test. F. Density of LDs formed on SLBs containing 10 mol% of the indicated lipid, 4 mol% DG, and DOPC. The SLBS were imaged 1hr after incubation with oleoyl-CoA. Colored squares show the mean of independent experiments; dots with the same color are quantifications of single visual fields used to calculate the means. Bars are the mean and std. dev. of 3 independent experiments; p value calculated from one-way ANOVA. G. *GAL1-DGA1* 3Δ *psd1*Δ *psd2*Δ cells were grown to mid-logarithmic growth phase in a medium with Etn, washed, resuspended in media with Cho and no Etn (left) or Etn and no Cho (right), and grown for 3 h. Estradiol was then added to the media to induce TG production, and cells were grown for 1 h. The cells were then stained with BODIPY and visualized. Bars = 5μm. H. Quantification of experiments shown in panel G. Gray dots indicate number of droplets in individual cells; colored squares are means of independent experiments with at least 500 cells per condition. Bars and lines show the mean and std. dev. of the three experiments; p value calculated from Welch’s t-test. I. Density of LDs formed on SLBs in assays with the indicated amount of PE, 4 mol% DG, and DOPC. The results are depicted as described in panel F.

We found the velocity of TG production also affects the number of LD formed in cells. For these experiments, we used yeast, which has two proteins that produce TG (Dga1 and Lro1) and two that produce primarily steryl esters (Are1 and Are2). Cells lacking all four proteins do not produce neutral lipids and lack LDs^50^. We used a strain that expresses *DGA1* under an estradiol regulatable *GAL1* promoter and does not express the other three neutral lipid synthesizing enzymes (*GAL1-DGA1* 3Δ). When grown in a medium without estradiol, this strain lacks LDs but begins to form them when estradiol is added to the medium. We found that the strain produces TG at different rates when different amounts of estradiol were added to the growth medium (Fig. S2E, F). Cells grown for 1 hour with 500 nM estradiol produced about as much TG as cells grown with 5 nM estradiol for 2 hours (Fig. S2E). The cells grown with 500 nM estradiol produced more LDs than cells grown with 5 nM estradiol despite producing similar amounts of TG (Fig. S2F). This indicates that in cells, as in artificial membranes, producing TG at a higher rate results in more LD formation. It should be noted that the yeast strain we used expressed seipin and the role of seipin in LD formation is addressed in subsequent sections. Collectively, these experiments reveal that LD nucleation rate is proportional to the velocity of TG production in membranes.

### ER-like phospholipid composition promotes LD nucleation

Recent studies^19,21,51^ indicate the phospholipid composition of the ER promotes TG de-mixing. To test this, we compared LD formation in SLBs made with an ER-like phospholipid mix^52^ (ER-mix; Fig S3A) or DOPC. When reactions were allowed to run to completion, the ER-mix SLBs formed 10-fold more LDs than DOPC SLBs (Fig. 3E), suggesting the lipid composition of the ER presents a lower energy barrier for TG phase separation than control DOPC membranes.

To dissect how the ER-like lipid mix promotes LD nucleation, we investigated the contribution of phospholipid type and acyl chain composition. SLBs composed of PCs with acyl chains that have a range of packing defects had no significant effect on LD production (Fig. S3B). In contrast, we found that the intrinsic curvature of phospholipids, which is related to the lipid’s effective molecular shape, had a significant effect on nascent LD formation. Phospholipids with little or no intrinsic curvature (cylindrical lipids, phosphatidylcholine (PC), phosphatidylserine (PS) and phosphatidylinositol (PI)) had no effect on nascent LD nucleation (Fig. 3F). In contrast, cone-shaped lipids phosphatidylethanolamine (PE) and phosphatidic acid (PA), and the inverted cone-shaped lipid lysoPC increased nucleation (Fig. 3F). We quantified TG synthesis and ruled out whether these lipids promoted LD nucleation by increasing sDGAT activity (Fig. S3C).

Following LD formation in real-time allowed a more comprehensive determination of the effects of cone and inverted-cone shaped lipids. When membranes contained either PE or lysoPC, LD nucleation time decreased (Fig. S3D, E). Even though PE and lysoPC have opposite molecular shapes, using membranes containing both lipids together also decreased LD nucleation time (Fig. S3D, E). It is likely that all non-cylindrical lipids decrease the energy barrier to LD nucleation^19,21,51^. Interestingly, non-cylindrical lipids have little or no effect on the growth rate of LDs after nucleation (Fig. S3F).

To test whether non-cylindrical lipids affect LD nucleation in cells, we used the yeast strain that allows us to measure LD nucleation (*GAL1-DGA1* 3Δ) and genetically modified it to allow us to control the amount of PE in the cells. It has been found that yeast lacking the PE synthesizing enzymes Psd1 and Psd2 have low PE when grown with exogenous choline and high PE when grown with ethanolamine^53^, a finding we confirmed (Fig. S3 G, H). We quantified LD formation in cells with either low or high PE and found that cells with low PE produced fewer LDs than cells with high PE (Fig. 3 G, H). It should be noted that cells grown with ethanolamine produced about 1.3-fold more TG than those grown with choline (Fig. S3I), but this is not enough to account for the 2-fold increase in LDs found in cells grown with ethanolamine (Fig. 3 G, H). Consistent with our results in cells, SLBs with 10 mol% PE produced ∼7-fold fewer LDs than those with 30 mol% PE (Fig. 3I). Altogether, our findings suggest that the presence of non-cylindrical lipids in the ER significantly increases the propensity of new LDs to form in the ER, probably by lowering the barrier to TG phase separation.

### Seipin accelerates LD nucleation

TG phase separation occurs once its concentration exceeds the critical concentration. Hence, we sought to determine this critical concentration in our LD formation assay. Since our assay relies on TG synthesis from DG, we estimated this concentration by comparing whether LDs formed in DOPC SLBs containing varying amounts of DG. SLBs were incubated with 1µM sDGAT and reactions were allowed to run to completion (1hr). The number of LDs formed was proportional to the starting DG concentration in the SLBs (Fig. 4A). No LDs were observed in membranes containing 1 mol% DG. With 2 mol% DG, few LDs occasionally form in the membrane (Fig. 4B). This indicates that the critical concentration for TG in these SLBs is approximately 2 mol%, in agreement with previous estimates^2,3,5,18,20^.

**Figure 4.**
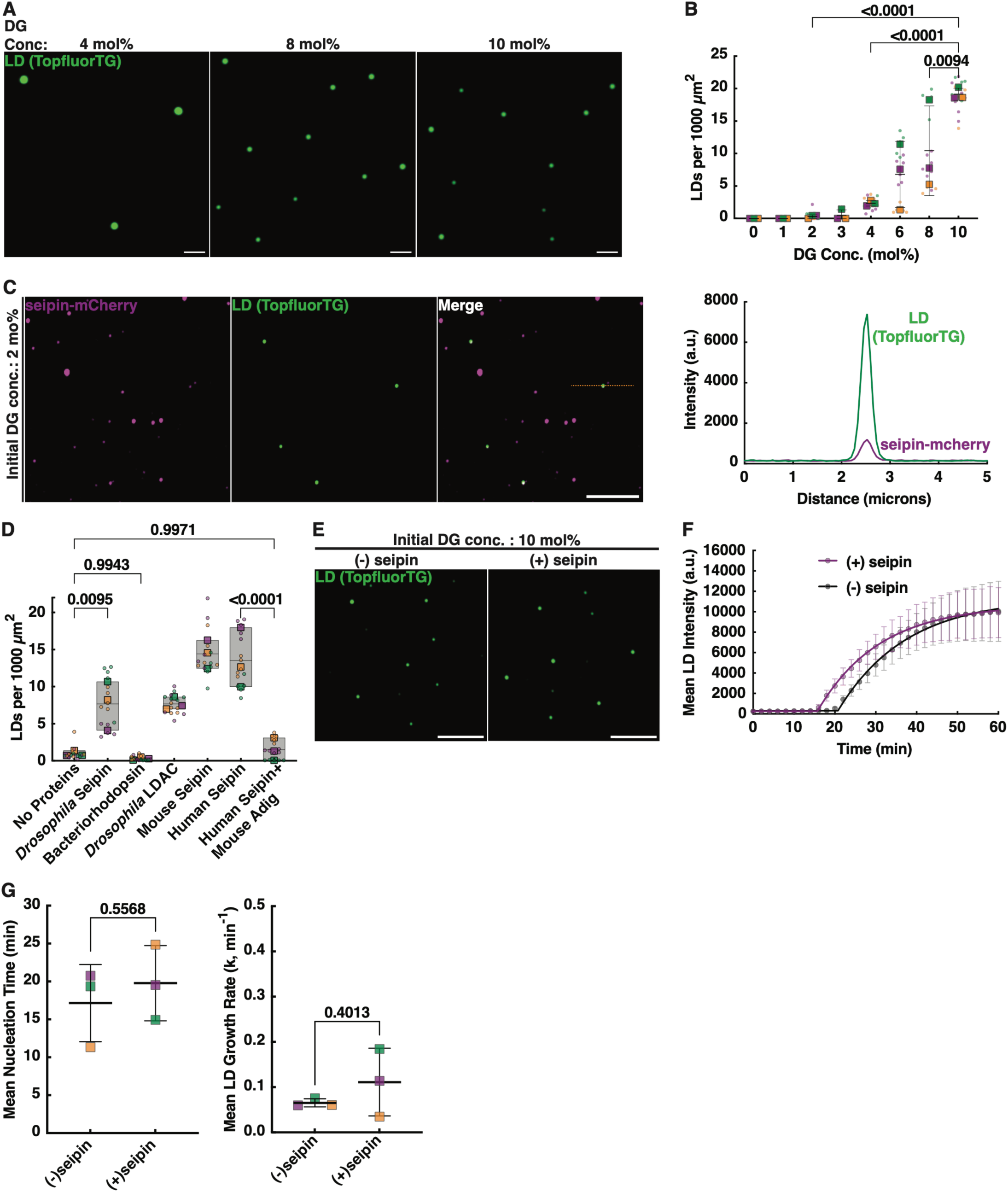
Seipin facilitates nascent LD nucleation. A. Images of SLBs from LD formation reactions performed as in Fig. 3A except all the reactions had 1000 nM sDGAT and the indicated amount of DG. Images taken 1 h after oleoyl-CoA addition. Bars = 2 μm. B. Density of LDs formed on SLBs in assays as in panel A. Colored squares show the mean of independent experiments; dots with the same color are quantifications of single visual fields used to calculate the means. Lines indicate the mean and std. dev. of the three experiments; p values calculated from one-way ANOVAs. C. LD formation assays were performed using SLBs with DOPC:DG:TopFluorTG (97.9:2:0.1 mol%) that contain seipin-mCherry. The SLBs were imaged 1 hour after oleoyl-CoA addition (left). Bar = 5µm. Fluorescence signal along dotted line (right). D. Lipid droplet formation assays were performed as in panel C except the SLBs were formed from proteoliposomes containing the indicated proteins. Box plots show mean, minimum and maximum of 3 independent experiments. Large colored squares show mean of individual experiments; dots with the same color are quantifications of single visual fields used to calculate the means; p values calculated from one-way ANOVAs. E. Lipid droplet formation assays were performed with SLBs that had an initial concentration of 10 mol% DG and that did or did not contain *Drosophila* seipin. Images acquired every 2 min. Panels show LDs formed 1 hour after oleoyl-CoA addition. Bar = 5µm. F. Quantification of experiments in panel E. Mean +/− std. dev. of LDs formed without (153 LDs) or with *Drosophila* seipin (90 LDs). G. Mean nucleation time (left) and growth rate (right) of LDs formed in assays as in panels E and F. Mean and std dev. of 3 independent experiments. p value calculated from Welch’s t-test.

While LDs appear to form stochastically in our assay, in cells they appear to form at seipin containing sites in the ER^26,39^. MD simulations and reconstitution of seipin in membranes with TG suggest TG molecules cluster within seipin^35,36^. Recent studies suggest LDAF1 might also help facilitate nucleation^18,39^. To investigate the role of seipin and its partner proteins in LD formation, we purified recombinant full-length *Drosophila melanogaster* seipin fused to mCherry at its C-terminus (seipin-mCherry). Size exclusion chromatography was employed to purify the oligomer as described previously^30^ (Fig. S4 A, B) and the oligomer was used to form proteoliposomes (scheme in Fig. S4C). The incorporation of seipin into proteoliposomes was verified by flotation and carbonate extraction (Fig. S4C-E). Seipin-proteoliposomes containing 2 mol% DG were used to generate SLBs and examined in our LD formation assay. We observed that seipin accelerates LD formation; all LDs formed colocalized with seipin (Fig. 4C). As a control, we showed that another multi-pass integral membrane protein, *Halobacterium salinarum* bacteriorhodopsin, does not support LD formation (Fig. 4D). Our findings reveal that seipin alone is sufficient to facilitate TG clustering and consequently accelerate LD nucleation.

Since LDAF1 has been suggested to be required for seipin to efficiently assemble LDs^18,39^, we next investigated whether LDAF1 further enhances LD nucleation. We purified *Drosophila melanogaster* LDAC (seipin + LDAF1) and used it to generate SLBs containing LDAC. Surprisingly we observed no significant increase in LD formation (Fig. 4D). One explanation for this result could be that seipin from *Drosophila* melanogaster may function independently of LDAF1. However, when we made SLBs that contained human or mouse seipin lacking LDAF1, they also support LD formation (Fig. 4D). Therefore, in our in vitro assay, LDAF1 is not required to nucleate LDs.

Recently, the microprotein Adig was shown to form a complex with seipin, and MD simulations suggested that Adig-rich seipin suppressed TG nucleation^41^. Therefore, using our LD formation assay, we determined whether the ability of seipin to form LDs was inhibited by Adig. When seipin-Adig complexes were incorporated into SLBs, seipin no longer supports LD formation. We used floatation to confirm that the Adig did not affect seipin incorporation into proteoliposomes (Fig. S4F). We conclude that Adig inhibits the ability of seipin to nucleate LDs in our assay, confirming the claim that it blocks seipin-mediated LD formation in cells^41^.

Besides nucleation, seipin has been implicated in other steps of LD biogenesis: droplet growth and the emergence of LDs from the ER membrane^31,32,37,40^. Our assay enables quantitative analysis of LD growth, thus providing a means to measure the role of seipin in LD growth after nucleation. To test this, we induced TG production in SLBs containing 10 mol% DG, either with or without seipin. We found there was no significant difference in the LD growth rate (Fig. 4E-G). Taken together, these experiments suggest that seipin initiates LD nucleation but has no effect on subsequent TG accumulation.

### PLIN3 stabilizes TG in nascent droplets, slowing Ostwald ripening

PLIN3 is one of the first proteins to arrive at forming droplets other than seipin and LDAF1^39^. PLIN3 binding to LDs is thought to be driven by its ability to sense packing defects on the surface LDs^54–56^ and PLIN3 binding to LDs has been found to reduce droplet surface tension^57^. However, how these properties contribute to early steps of LD formation is not clear. To better understand it, we purified mCherry-XL labelled PLIN3 and carried out LD formation assays with and without PLIN3. PLIN3 accumulated on nascent LDs, as it does in cells (Fig. S5A), but it did not promote nucleation (Fig. S5B). Tracking LD populations over time revealed that LDs formed in the presence of PLIN3 were more resistant to dissolution by Ostwald ripening than those in control reactions lacking PLIN3 (Fig. 5A). Comparing the mean intensity of shrinking droplets, PLIN3 slowed dissolution of LDs (Fig. 5B). Consistent with this, we observed PLIN3 coated droplets had slower TopFluorTG recovery after FRAP compared to control LDs (Fig. S5C), indicating PLIN3 binding reduces TG mobility in droplets and consequently reduces Ostwald ripening.

**Figure 5.**
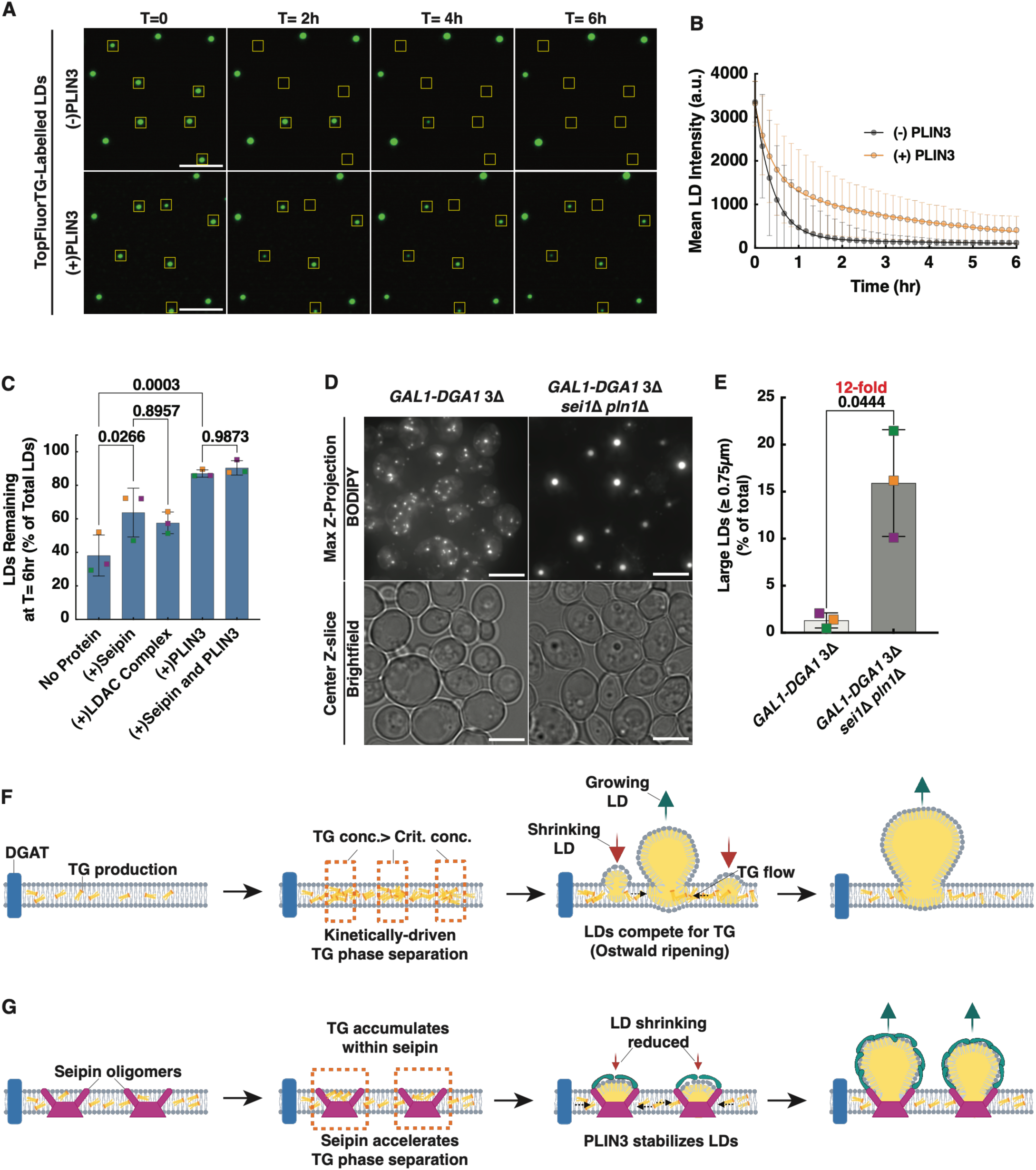
PLIN3 stabilizes TG in nascent droplets, slowing Ostwald ripening. A. LD formation assays were performed using SLBs containing 10 mol% DG and 25nM sDGAT, either with or without PLIN3. LDs were allowed to form for 30 min and then images were acquired every 10 minutes for 6 hours. Yellow boxes indicate shrinking LDs. Bar = 5µm. B. Quantification of mean fluorescence intensity of shrinking LDs from experiments as in panel A; mean and std. dev. of at least 450 LDs per condition. C. LD formation assays were performed on SLBs containing DOPC:DG:TopFluorTG (97.9:2:0.1 mol%) and the indicated proteins. Bars show percent of LDs that remained at T= 6hr. Colored squares show the mean of independent experiments. Lines indicate the mean and std. dev. of the three experiments; p values calculated from one-way ANOVAs. D. Cells were grown to mid-logarithmic growth phase, and estradiol was added to induce TG production. *GAL1-DGA1* 3D cells were grown for 1 h and *GAL1-DGA1* 3D *psd1D psd2D* cells for 2 hrs. The they were then stained with BODIPY and visualized. Bars = 5μm. E. Quantification of experiments shown in panel E. F. Scheme of LD formation after TG synthesis in a membrane. Schematic created using BioRender (BioRender.com). G. Scheme of LD formation after TG synthesis in a membrane containing seipin and PLIN3. Schematic created using BioRender (BioRender.com).

Seipin has also been shown reduce Ostwald ripening of LDs^26^. Therefore, we compared effects of seipin alone, LDAC, and PLIN3 on ripening in our LD formation assay. We confirmed the starting density of droplets for these ripening experiments was similar (Fig. S5D). In our assays too, seipin slows nascent LD ripening (Fig. 5D), perhaps by stabilizing junctions between the ER and LDs^26,33^. LDAC was not better than seipin alone at slowing ripening (Fig. S5D). However, PLIN3 was significantly better at slowing ripening alone or in combination with seipin.

We next determined whether the ability of seipin and perilipin to slow Ostwald ripening affects the formation of nascent LD in cells. For this, we used the *GAL1-DGA1 3*Δ yeast strain and an isogenic strain also lacking seipin and Pln1, the sole perilipin in yeast (*GAL1-DGA1 3*Δ *sei1Δ pln1Δ*). In these strains, LD formation begins when estradiol is added to the medium to induce TG production. The strain lacking seipin and Pln1 produced about as much TG in 2 hours as the strain expressing these proteins made in 1 hour (Fig. S5E). Using these induction condition, the *GAL1-DGA1 3*Δ *sei1*Δ *pln1*Δ produced fewer and larger LDs (Fig. 5E, F) as previously observed^58^. We propose that in this strain, small nascent LDs are destabilized leading to increased ripening and the production of fewer, larger LDs. Collectively, our results reveal that perilipins stabilize nascent LDs, slowing the dissolution of small LDs by ripening.

## Discussion

Cells produce LDs in response to various metabolic stresses and require mechanisms that facilitate regulation of this process. We sought to understand how cells determine whether TG is directed to new droplet biogenesis or used to expand existing droplets. We reconstituted LD formation to assess the distribution of TG as it is produced in a membrane containing an ensemble of membraned-embedded LDs, mimicking what occurs in the ER. This revealed that membrane-embedded LDs suppress new LD formation by absorbing TG from the membrane and destabilizing nascent LDs by Ostwald ripening. Increasing the velocity of TG production, promoted TG demixing and therefore LD nucleation. Similarly, TG demixing is promoted in the ER by non-cylindrical phospholipids, probably by altering the LD-membrane interfacial tension as has been suggested^51,59^. The LD biogenesis proteins seipin and PLIN3 promote nascent LD formation at distinct steps by overcoming the suppression of LD formation by membrane-embedded LDs.

Together, these findings show how cells can channel TG towards (or away from) nascent LD biogenesis. In a membrane with only DGAT, TG will spontaneously phase separate. Our findings show that the velocity of TG production and membrane phospholipid composition affect TG nucleation, indicating the rate of TG phase separation events is kinetically determined (Fig. 5F). This implies LDs that nucleate first, grow larger by accumulating TG molecules from the membrane, which then leads to droplet coursing by Ostwald ripening (Fig. 5F). Seipin promotes LD nucleation and PLIN3 stabilizes nascent LDs (Fig. 5G). Other PLINs or LD-bind proteins could similarly stabilize nascent LD instead of PLIN3 during LD formation in some cell types or growth conditions.

We show seipin accelerates TG phase separation, controlling the kinetics of TG demixing and allowing LDs to grow spontaneously within seipin after nucleation. We show that seipin alone is sufficient to initiate this process, perhaps by simply clustering TG, which increases the probability of TG phase separation. Therefore, one of the main functions of seipin, beyond TG nucleation, could be to ensure that seipin is at all ER-LD junctions. In our assay, LDAF1 does not seem to affect LD nucleation, growth or ripening, raising the question of LDAF1’s role in LD formation. We also demonstrate that Adig bound seipin cannot nucleate TG, consistent with the function of Adig in mice^41^. Future work with seipin mutants in our assay will enable tracking the stages of LD formation. This will allow more mechanistic insights into how seipin promotes LD nucleation, organizes TG and other neutral lipids, stabilizes the ER-LD interface, and works together with partner proteins.

## Materials and Methods

### Materials

All reagents, unless otherwise indicated, were purchased from Sigma-Aldrich. All unlabeled and fluorescent lipids were from Avanti Polar Lipids (Croda), and radiolabeled lipids were from American Radiolabeled Chemicals. Full length wild-type *Halobacterium salinarum* bacteriorhodopsin was purchased from Cube Biotech (Germany).

### Methods

#### Construction of plasmids

All primers used for plasmid construction were purchased from Integrated DNA Technologies (IDT) and are listed in Table S1. sDGAT from *Acinetobacter baumannii* (obtained from American Type Culture Collection) was amplified and cloned into a pET15b vector with an N-terminal 6×His tag and a C-terminal Strep II tag. mCherry was fused at the C terminus to generate sDGAT–mCherry. A gBlock (IDT) encoding codon-optimized *PLIN3*, based on a previously published sequence ^54^, was amplified and cloned into pET15b with an N-terminal Strep II tag and a C-terminal 6×His tag. A TEV protease cleavage site was inserted between PLIN3 and the His tag to allow tag removal. pBAD-mCherry-XL (R125K) was a gift from Ralph Jimenez (Addgene plasmid #188054). PCR primers were used to amplify and insert mCherry-XL at the N terminus of PLIN3. *Drosophila* seipin-6xHis was a kind gift from the Walther and Farese lab. Using PCR primers, mCherry was fused at the C-terminus to generate seipin-6xHis-mCherry. pACYCDuet-1_Ec_rpoZ was a gift from Seth Darst (Addgene plasmid # 128837). A gBlock (IDT) encoding a codon optimized *Drosophila* LDAF1 with an N-terminal ALFA tag was amplified with PCR primers for insertion into the plasmid.

#### Expression and purification of proteins

sDGAT, PLIN3, Drosophila seipin and Drosophila LDAC complex constructs were purified by expression in *E. coli* BL21(DE3) (New England Biolabs, USA), grown in auto-induction medium (Formedium, UK) for 36–48 h at 18 °C. Pellets were stored at −80 °C. For purification of sDGAT and PLIN3 constructs, pellets were thawed, resuspended in 20 mM HEPES (pH 7.4), 500 mM NaCl, and lysed with a cell disruptor (Constant Systems). Lysates were clarified by centrifugation (20,000 × *g*) supernatants were bound to HisPur Cobalt resin (Thermo Scientific), washed with 20 mM HEPES (pH 7.4), 500 mM NaCl, and eluted with 20 mM HEPES (pH 7.4), 500 mM NaCl containing 200 mM imidazole. Eluates were applied to a 5mL Strep-Tactin column (Cytiva) and washed with 20 mM HEPES (pH 7.4), 150 mM KCl buffer. Proteins were eluted with 2.5 mM desthiobiotin in 20 mM HEPES (pH 7.4), 150 mM KCl buffer. The 6×His tag on PLIN3 was cleaved with His-tagged TEV protease. For Drosophila seipin and Drosophila LDAC complex purification, cells were transformed with either seipin containing plasmid alone or seipin and LDAF1 plasmids respectively and purified as described previously^30^. Human seipin, mouse seipin and human seipin with mouse Adig complexes were produced using Expi293 Expression System (Thermo Fisher Scientific, no. A14635) and purified as described previously^41^, except oligomers were separated using Superose 6 Increase column 10/300GL (Cat.no. GE290915-96). Purified proteins were stored at 4 °C before use. Before any microscopy and biochemical assay, purified proteins were centrifuged (120,000 × *g*, 30 min) to remove aggregates. Protein concentrations were measured using Pierce Detergent Compatible Bradford assay Kit (Thermo Scientific).

#### Liposome preparation

Lipids were mixed at the stated ratios in glass tubes, dried as a thin film under nitrogen, and placed under vacuum for 1 h to remove residual solvent. Films were hydrated with 500 µL Milli-Q water, incubated at 50 °C for 30 min, and extruded through 50-nm polycarbonate filters (Avanti Polar Lipids).

#### Supported lipid bilayer (SLB) preparation

All assays were carried out in assay buffer (20 mM HEPES, pH 7.4, 150 mM KCl). Time-lapse experiments were performed on SLBs prepared in flow chambers. Piranha-cleaned glass coverslips were assembled into FCS2 chambers (Bioptechs, USA). A mixture of 250 µM liposomes and 1 M NaCl was incubated on the coverslip for 30–45 min to allow spontaneous fusion and SLB formation. Chambers were washed sequentially with Milli-Q water and assay buffer containing 1 mg/mL BSA. Purified proteins were added at the indicated concentrations and incubated for 10 min. TG synthesis was initiated by addition of 18 µM oleoyl-CoA in assay buffer supplemented with 1 mg/mL BSA and an oxygen scavenger cocktail^60^. For LD distribution measurements, including experiments with proteoliposomes, SLBs were prepared as above on 4-well or 8-well glass-bottom chambers (ibidi, USA).

#### Fluorescence microscopy and image analysis of LDs on SLBs

For Fig. 1B, images were acquired with a 100×/1.4 NA oil-immersion objective on a DeltaVision Elite wide-field inverted epifluorescence microscope (Applied Precision, GE Healthcare) equipped with an environmental chamber, a 100×/1.4 NA objective (Olympus), and a pco edge 4.2 CLHS sCMOS camera (Excelitas, USA). Images were deconvolved using the “conserved ratio” method (softWoRx, Applied Precision).

All other SLB experiments were imaged with a 100×/1.4 NA oil-immersion objective on a Nikon Eclipse Ti2 confocal microscope equipped with a Yokogawa CSU-W1 spinning disk and an ORCA-Fusion BT sCMOS camera (Hamamatsu). Images were acquired with NIS Elements (Nikon) and analyzed using Fiji (version 1.54p, National Institutes of Health, ^61^) and NIS Elements (version 5.42.03). Statistical analyses were performed in Prism 10 for MacOS (GraphPad).

For time-lapse experiments, LD mean intensity was quantified from movies. Maximum intensity projections were generated, thresholded to create binary masks of individual droplets, and overlaid on the original images. Droplet intensities per frame were measured using Fiji’s Plot Z-axis Profile plugin. Mean droplet intensity over time was plotted in GraphPad Prism and fit to a plateau followed by a one-phase association equation to calculate mean nucleation time and mean growth rate.

For Fig. S1E, mean intensity and area of individual droplets from multiple images were quantified using Fiji’s Particle Analysis plugin. Values were plotted in GraphPad Prism and fit to a centered second-order polynomial equation.

For 6 h shrinkage experiments, images of the same field of view were aligned using Fiji’s Linear Alignment by SIFT plugin prior to quantification with the Plot Z-axis Profile plugin. For Fig. 5B and 5E, LD intensities over time were plotted in GraphPad Prism and fit to a two-phase decay equation.

FRAP experiments were carried out on the Nikon spinning-disk confocal microscope using a 405-nm diode laser. Mean intensities of bleached and reference ROIs were quantified with the Time Measurement tool in NIS Elements and normalized to pre-bleach values.

#### Electron microscopy of LDs

SLBs and LDs were prepared on 12-mm glass coverslips (Cat. no. 26023, Ted Pella, USA) as described above. Samples were fixed with 2.5% glutaraldehyde (Electron Microscopy Sciences, USA) for 2 h, rinsed twice with 0.1 M cacodylate buffer (pH 7.4), and post-fixed with 2% osmium tetroxide in 0.1 M cacodylate buffer for 2 h. Fixed samples were washed with double-distilled water, dehydrated through a graded ethanol series (10–100%), and dried by critical point drying (SAMDRI-795, Tousimis; purge time 20 min). Samples were sputter-coated with a gold–palladium mix and mounted on SEM stubs for imaging.

All samples were imaged on a Zeiss GeminiSEM at low magnification (∼4800×, 25 kV) and high magnification (∼20,500×, 25 kV). Droplet diameters were measured from calibrated images using Fiji.

#### Giant Unilamellar Vesicle (GUV) preparation

Chloroform stock solutions of 1,2-dioleoyl-sn-glycero-3-phosphocholine (DOPC, 18:1), diacylglycerol (DAG 18:1), Ni–NTA–lipid, TopFluor TG (TG 18:1,C11), and rhodamine–DOPE (Avanti Polar Lipids, Croda) were mixed at a molar ratio of 93.3:4:2.5:0.1:0.1 to yield a 2 mM lipid mixture. 30–50 μL of this mixture were deposited onto indium tin oxide (ITO)–coated glass slides, dried under a gentle nitrogen stream, and desiccated under vacuum for 1 h. Lipid films were rehydrated with a sucrose solution (300 ± 10 mOsm) and subjected to an alternating current electric field (1.4–1.6 Vpp, 100 Hz) at 37 °C for 2 h to promote Giant Unilamellar Vesicle (GUV) formation.

#### Preparation of GUV Samples for Microscopy

GUVs were imaged in 4-well glass-bottom plates (Ibidi). Wells were passivated with 1 mg/mL bovine serum albumin (BSA) for 10 min, rinsed three times with assay buffer (20 mM HEPES, pH 7.4, 150 mM KCl), and GUVs were deposited in the chamber sDGAT and 1 mg/mL BSA was prepared, gently mixed, and added to the imaging chamber. The samples were incubated for 10 min, after which 18 μM oleoyl–CoA in assay buffer was added immediately before initiating time-lapse imaging.

#### Microscopy and quantification of GUVs

GUVs were imaged at 30 °C using a spinning disk confocal microscope with a 60× air objective. Images were acquired every 5 min for 0.5 hr. Fluorescence intensity was measured using Fiji.

#### LD formation in COS-7 cells

COS-7 cells were seeded in 4-well glass-bottom chambers (ibidi, USA) and cultured for 18 h in DMEM without FBS (Thermo Scientific, USA). Cells were incubated with BODIPY 493/503 (Thermo Scientific, USA) at a final concentration of 1 µg/mL for 20 min and washed twice with DMEM lacking phenol red (Thermo Scientific, USA) and FBS before imaging.

Time-lapse images of LD formation were acquired on a Nikon Ti2-E motorized inverted widefield fluorescence microscope (Nikon) equipped with a triggered PInano P-736 piezo Z stage for z-axis control (Physik Instrumente LP), a SPECTRA III 8-line LED light source (Lumencor), a CFI60 Plan Apochromat Lambda D 60×/1.42 NA oil-immersion objective (Nikon), a Kinetix 22 back-illuminated sCMOS camera (Teledyne Photometrics), and an Okolab black enclosure maintained at 37 °C (Okolab). The system was controlled with NIS-Elements software (Nikon). Nascent LD formation was initiated by adding 400 µM oleic acid (OA) in BSA (from a filtered stock containing 5mM OA 15%w/v BSA), and z-stack images of the cells were acquired every 20 s for 35 min.

For Fig. 1G and Movie 3, images were first deconvolved using the Richardson–Lucy method. Maximum intensity projections were then generated for each time point, and LUTs were adjusted in NIS-Elements software (AR 5.42.03, Nikon).

#### Quantification of LD formation in COS-7 cells

Maximum intensity projections were generated in FIJI from the raw images for each time point and converted into Imaris-compatible files. Nascent LD formation was quantified over time using the standard *Spot Detection* tool with a 0.3 µm size setting to identify LDs and tracked in Imaris (11.0.1, Oxford Instruments).

#### Saccharomyces cerevisiae culture

Yeast strains were engineered from the parental *are1Δ are2Δ lro1Δ GAL1-DGA1* pAGL strain (gift from Jonathan Friedman, UT Southwestern Medical Center) using established methods and primers listed in Table S1. All strains used in this study are listed in Table S2.

Cells were grown in synthetic complete (SC) medium containing 2% glucose, 0.67% yeast nitrogen base (Becton Dickinson), and amino acid drop-out mix (US Biological) at 30 °C. Estradiol was added at the stated concentrations to mid-logarithmic phase cultures to induce Dga1 expression.

For experiments investigating the role of PE on LD formation in cells, precultures were grown in SC medium with 4 mM Etn. Cells were washed twice in SC medium, diluted to OD_600_ = 0.4, and split into two cultures containing either 4 mM Etn or 4 mM Cho. After 4 h, TG production was induced by addition of 50 nM estradiol to these mid-log phase cultures for 1 h before imaging.

#### Yeast cell microscopy

Live yeast cells were imaged at 30 °C on a Nikon Eclipse Ti2 wide-field microscope with a 100×/1.4 NA objective (Nikon), an OKOlab chamber, a KinetiX 22 sCMOS camera (Teledyne Photometrics), and a Spectra III 8-line LED light source (Lumencor). For all images, Z-stacks (fluorescence and brightfield) were acquired over 5 µm with 0.2 µm steps. Images acquired and contrast adjusted and denoised with NIS Elements (version 5.42.03, Nikon).

#### Quantification of LD numbers per cell

Images were opened in Fiji (v1.54p National Institutes of Health) and cell contours were segmented using the Cellpose3 plugin (Janelia Research Campus, ^62^. Images were converted to Imaris-compatible format and imported into Imaris (Oxford Instruments). Images were deconvolved, and LDs were detected using the *Spot Detection* tool with a 0.3 µm size setting; the quality threshold was adjusted to optimize detection. The Cellpose3-generated contour masks were overlaid, and Imaris quantified the number of LDs per cell. Data were plotted and analyzed in GraphPad Prism.

#### Quantification of TG

Overnight yeast precultures were diluted into fresh medium, adjusted to equal optical density, and grown to mid-logarithmic growth phase before addition of estradiol at the indicated concentrations. At each time point in the experiment, 25 mL of culture was withdrawn, and metabolism was arrested with addition of 10 mM NaN₃ and NaF and incubation for 10 mins on ice. Cells were pelleted and stored at −80 °C.

Pellets were thawed on ice, resuspended in 1 mL ice-cold Milli-Q water, and transferred to 2 mL screw-cap tubes. Cells were pelleted at 4 °C, mixed with 0.3 mL glass beads, and resuspended in 0.3 mL ice-cold Milli-Q water. Lysis was performed in a Precellys24 homogenizer (Bertin) with three cycles of 30 s at 5500 rpm and 60 s pauses between cycles, at 4 °C. Lysates were extracted twice with 0.6 mL water, and pooled. Protein concentration was determined by Bradford assay (Pierce) using 5 µL of lysate.

For lipid extraction, 1 mL lysate was mixed with 6 mL MeOH:CHCl₃ (2:1) and 0.6 mL water. Phase separation was induced by addition of 2 mL 0.9% NaCl (VWR) and the CHCl_3_ phase was back extracted with 2 mL 2 M KCl. After centrifugation (1500 × g, 5 min), the lower phase was transferred to fresh glass tubes and dried under a gentle nitrogen flow. Samples were resuspended in 200 µL hexane, and 100 µL was injected into an Agilent 1260 Infinity II HPLC system (Agilent Technologies) equipped with a Zorbax CN column (5 µm × 150 mm, Agilent). TG was separated with hexane:tert-butyl methyl ether (97:3) run for 3 min, followed by a gradient to 80:20 run over 7 min, and then held for 5 min. TG was detected by evaporative light scattering (ELSD, Agilent Technologies), and peak areas were quantified. A standard curve generated with pure triolein (Avanti Polar Lipids, Croda) was used to convert peak area to nmol TG amounts. TG levels were normalized to total protein per time point for each sample.

#### Phospholipid quantification

Precultures were grown in SC medium with 4 mM Etn. Cells were washed twice in SC medium, diluted to OD_600nm_ = 0.4, and split into two 10 ml cultures containing either 4 mM Etn or 4 mM Cho. 50 µCi [³H]palmitic acid (ART0129A, American Radiolabeled Chemicals) was added to these cultures and grown for 4 h at 30 °C. TG production was induced by addition of 50 nM estradiol for 1 h. Cells were pelleted and stored at −20 °C.

Lipids were extracted using methods described above. Dried lipid films were resuspended in 30 µL CHCl₃:MeOH (2:1) and loaded onto Silica thin layer chromatography (TLC) plates (Merck). Plates were developed twice for 2 h each in CHCl₃:EtOH:H₂O:triethanolamine (30:35:7:35) and air-dried. Radioactive lipids were quantified with a RITA Star TLC analyzer (Raytest). Phospholipid identity (PC, PI, PS, PE) was determined using cold standards run in parallel lanes and revealed by copper sulfate staining. Radioactivity in each phospholipid band was quantified and normalized to the total signal from the four phospholipids.

#### Measurement of DGAT activity

Stocks of liposomes (2 mM) were prepared that contained DOPC:DG:DOGS-Ni-NTA,TopFluorTG:indicated lipid (83.4:4:2.5:0.1:10 mol %) in assay buffer (20 mM HEPES, pH 7.4, 150 mM KCl). The liposomes also contained 50 mCi/mL [^3^H]DG (ART 2185, American Radiolabeled Chemicals). Reactions containing 250 mM liposomes and 1 mM sDGAT in assay buffer and were started by addition of 18 mM oleoyl-CoA. At the indicated times, 1 mL aliquots were withdrawn, and lipids were extracted and separated on a silica TLC plates developed with hexane:diethyl ether:acetate (70:30:1). The amount of [^3^H]TG formed was determined with a RITA Star TLC analyzer (Raytest).

#### Monte Carlo Simulations

The free energy of formation of an LD composed of *n* TG molecules out of a reservoir of TG dispersed in the surrounding membrane is given by:

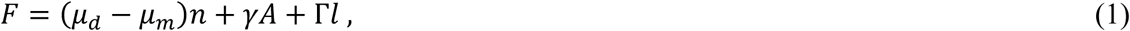

where *μ_d_* and *μ_m_* are the chemical potentials of TG in the droplet and in the reservoir, respectively, *γ* is the LD’s surface tension, *A* is the area of the LD’s surface, Γ and *l* are, respectively, the line tension and the length of the contact line formed between the LD base and the membrane.

The area, *A*, and contact line length, *l*, depend on the shape of the LD. The latter was modeled as a truncated sphere of radius *R* with a wetting angle *θ*. We assume the wetting angle to be independent of the radius of the sphere, and depends only on the substrate properties and LD surface tension, *γ*. The volume of an LD is proportional to the number of molecules in the droplet, *n*. This gives the following relation between the radius of the droplet and the number of molecules inside it:

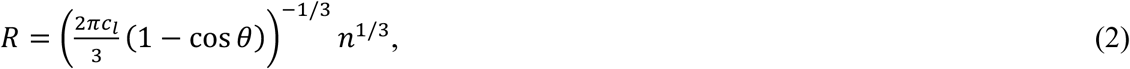

where *c_l_* is the density of TG molecules in the droplet.

The difference in chemical potential, appearing in Eq. (1), is a function of the TG concentration in the membrane reservoir, *c*, and can be written as

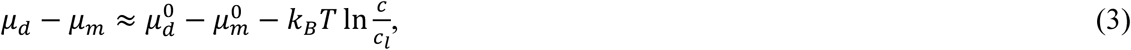

where 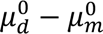 is the concentration-independent contribution.

Plugging Eqs. (2-3) into Eq. (1), together with the explicit expressions for surface area and contact length, gives for the LD’s free energy:

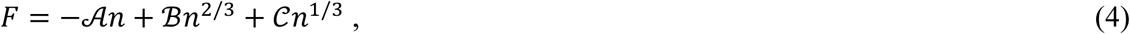

where 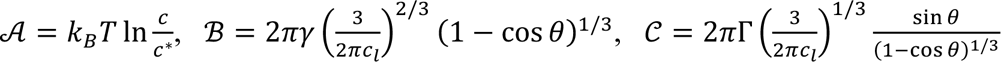, and 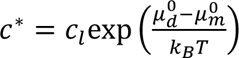. The dependence of the energy (Eq. 4) on the number of molecules comprising a LD, is described by a bell-like curve with a maximum at the critical number of molecules, *n*^∗^. According to this, a stable LD requires an over-critical number of molecules, *n* > *n*^∗^. Droplets with sub-critical sizes, *n* < *n*^∗^, are predicted to shrink and disappear. The energy at the critical number, *F*(*n*^∗^), represent an energy barrier the system has to overcome to form a stable LD. The energy barrier can be found by the relation 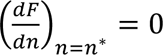, which gives:

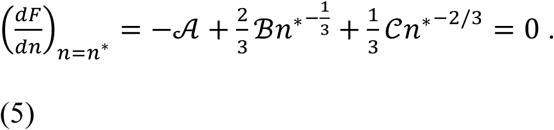

The meaningful solution of (Eq.5) is given by:

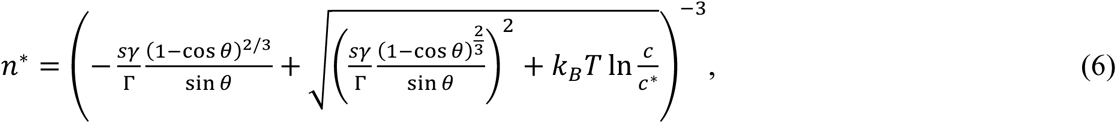

where 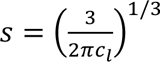. Plugging Eq. (6) into Eq. (4), we obtain an expression for the energy barrier, *F*^∗^, for forming a stable LD:

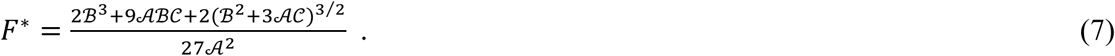

An LD grows by absorbing TG molecules from the membrane reservoir through the contact. The growth rate is proportional to the difference in the TG chemical potentials in the LD and the reservoir. Therefore, the time dependence of the LD’s number of molecules is determined by:

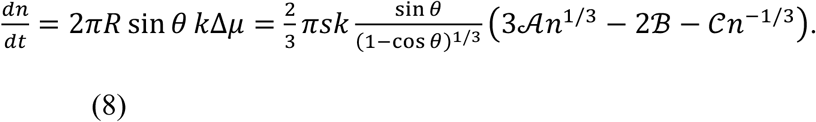

where *k* is a kinetic parameter, independent of the LD size.

We performed Monte Carlo simulation of the dynamics of redistribution of TG molecules between the membrane reservoirs. We constructed a square grid of potential LD nucleation sites imposed on a membrane patch. TG molecules were assumed to be produced everywhere along the membrane uniformly at a constant rate, *V_p_*. In our sumulation, at each time step, d*t*, each potential nucleation site made *f* binary attempts to become a critically sized LD. The overall success probability of a potential nucleation site (subrcritical LD) to become a critically-sized LD, during a time step d*t*, was taken as:

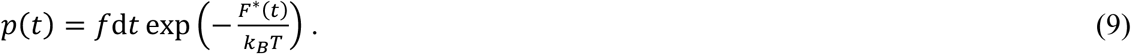

During each time step, any existing LD absorbed TG molecules from the bulk at a rate given by Eq. (8). At the end of a time step, we update: (i) the number of formed LD (ii) the size of each LD, and (iii) the concentration of bulk TG according to LD formation and growth, and the TG production rate. The simulation recorded the number of TG molecules per unit area of the membrane reservoir over time, and ended when the formed LD molecules depleted the TG concentration in the membrane to a value below the critical concentration. For every data point produced, the simulation was repeated three times to record the average and standard deviation. When varying the surface tension, we concurrently varied the wetting angle *θ* according to Young’s relation, cos *θ* ∼*γ*^−1^.

#### Proteoliposomes preparation

Before incorporation, bacteriorhodopsin was labelled by adding equimolar amounts of Atto 565 NHS ester, according to the manufacturer’s instructions. Lipids were aliquoted at the stated ratios in glass tubes, dried under nitrogen, and placed under vacuum overnight. Dried lipid films were hydrated with 1mL of buffer containing 20mM HEPES, ph7.4, 150mM KCl, 1% (w/v) Triton-X-100 and 0.05% (w/v) DDM to form detergent destabilized liposomes^63,64^. 16 ug of proteins were added to these destabilized liposomes at a 1:250 (w/w) protein to lipid ratio. The mix was incubated at room temp for 1h. Detergent was removed by successive addition of SM-2 Biobeads (40mg for 1h at RT, 40 mg for 2h at RT and 120mg overnight at 4°C). Protein incorporation was confirmed using sucrose density gradient centrifugation. Proteoliposomes were used immediately or stored at 4°C.

#### Sucrose density gradient centrifugation

All gradients were prepared on ice. Proteoliposomes were mixed with an equal volume of 60% (w/v) sucrose in 20 mM HEPES pH 7.4, 150 mM KCl. Gradient was formed by overlaying this mixture with 25% (w/v) sucrose in the same buffer, followed by buffer without sucrose. Ultracentrifugation was performed in a TLS-55 swinging-bucket rotor (Beckman) at 200,000 × g for 3h at 4°C. Equal-volume fractions were then withdrawn from top to bottom, diluted 4-fold with buffer without sucrose, and centrifuged again at 150,000 × g for 1 h at 4 °C to pellet the proteins.

For Fig. S4E, *Drosophila* seipin–mCherry proteoliposomes were mixed with an equal volume of 60% (w/v) sucrose and 200 mM sodium carbonate and incubated on ice for 30 min and fractionated using sucrose density gradient centrifugation and analyzed by western blotting.

#### Western blotting of gradient fractions

Pellets prepared from gradient fractions were resuspended in 2× NuPAGE LDS sample buffer (Thermo Fisher Scientific). Samples were resolved using 4-12% NuPAGE Bis-Tris gels (Thermo Fisher Scientific) and then transferred to nitrocellulose membranes (Bio-Rad). Membranes were blocked with 3% BSA for 1h at room temperature and incubated with primary antibodies (rabbit polyclonal Anti-mCherry, Cat. No. 600-401-P16, Rockland Inc.) overnight at 4°C. Membranes were washed and incubated with secondary antibodies (IRDye 800CW Donkey Anti-Rabbit IgG, Li-COR) for 2h at RT. Membranes were washed and imaged using Odyssey scanner (LI-COR).

#### Dot blot analysis of gradient fractions

Pellets prepared from gradient fractions were resuspend in 50ul of 20mM HEPES pH 7.4, 150mM KCl 5mM sodium azide and applied to a nitrocellulose membrane assembled in the dot blot apparatus (Bio-Rad) as per the manufacture’s protocols. After 1h incubation, the membrane was blocked by adding 5% BSA for 1h at room temperature and then incubated with primary antibodies overnight at 4 °C. The membrane was then washed and incubated with secondary antibodies for 2h at RT and imaged using ChemiDoc MP Imaging system (BioRad). Primary antibodies were Rabbit Anti-Strep-Tag-II (ab76949, Abcam) for human seipin and rabbit polyclonal anti-mCherry (Cat. No. 600-401-P16, Rockland Inc.) for *Drosophila* seipin. Secondary antibody was IRDye 800CW Donkey Anti-Rabbit IgG, LI-COR. Atto-565 labeled-bacteriorhodopsin was detected using the rhodamine filter.

#### Statistical Analysis

All statistical analysis unless otherwise mentioned was carried out using Graphpad Prism. For statistical comparison, when two mean values were compared against each other, a Welch’s *t*-test was conducted in GraphPad Prism to calculate and plot *P* values in the graph. The *P* values are indicated in the figures. To compare multiple mean values against each other a one-way analysis of variance (ANOVA) was used. The *P* values are indicated in the figures. Data used in this study are available in the article. No statistical method was used to predetermine sample size. No data were excluded from the analyses, and no randomization was performed in this study. Data distribution was assumed to be normal, but this was not formally tested.

## Acknowledgments

We thank Drs. Kate Luby-Phelps, Khuloud Jaqaman, and Marcel Mettlen for assistance with image analysis, Jonathan Friedman and Joel Goodman for yeast strains, and Mike Henne, Thomas Pucadyil for helpful discussions and comments on the manuscript. We thank the Phoebe Doss and Ethan Ozment for assistance with SEM.

## Author contributions

Conceptualization: RD, WAP and MMK

Methodology: RD, AT, AB, MMK, EH, CL, PES

Investigation: RD, AT, AB, EG, CL, EH, DB

Funding acquisition: WAP

Writing – original draft: RD and WAP

## Data and materials availability

All data are available in the main text or the supplementary materials. For correspondence

---

**Table S1.**
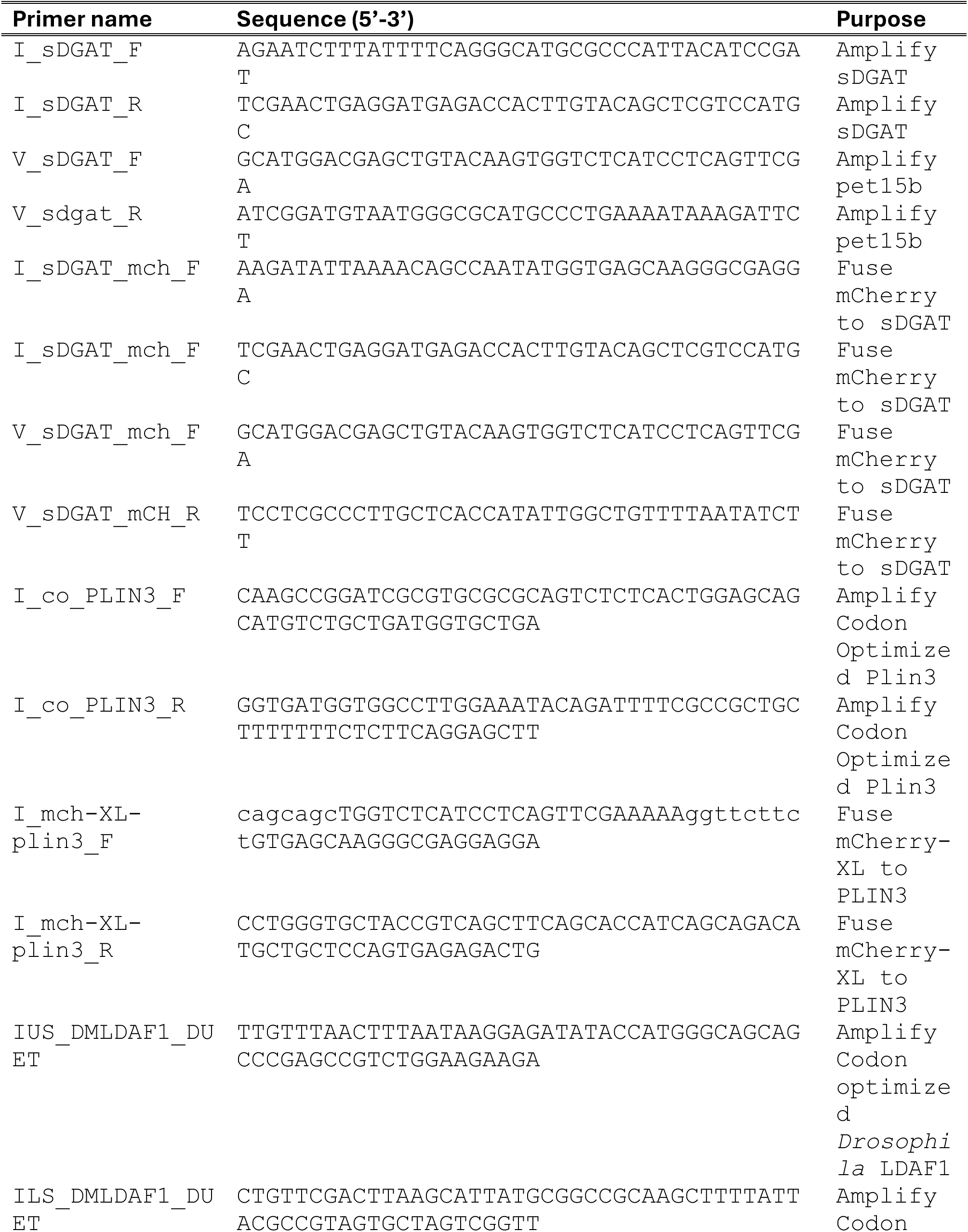
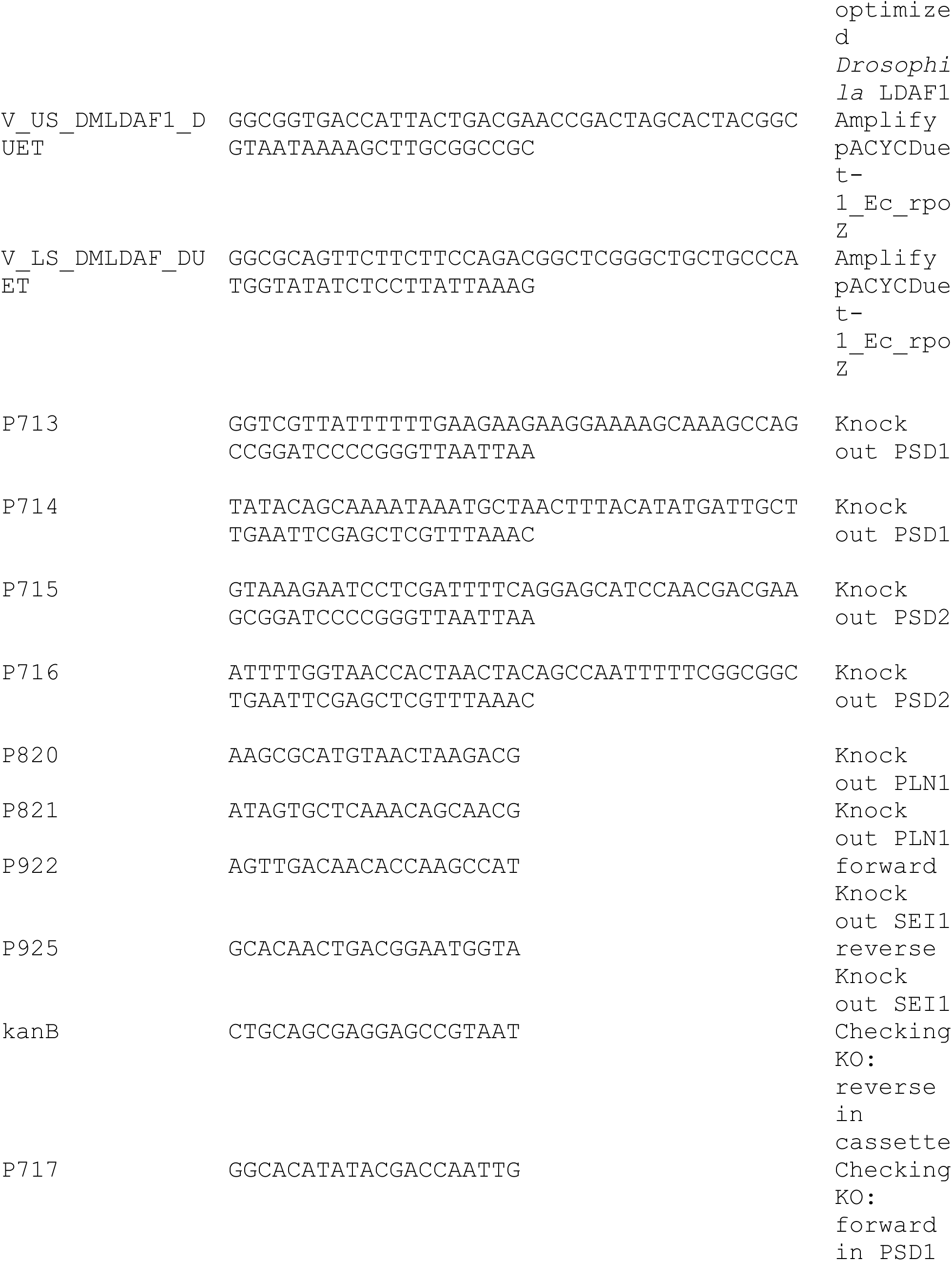
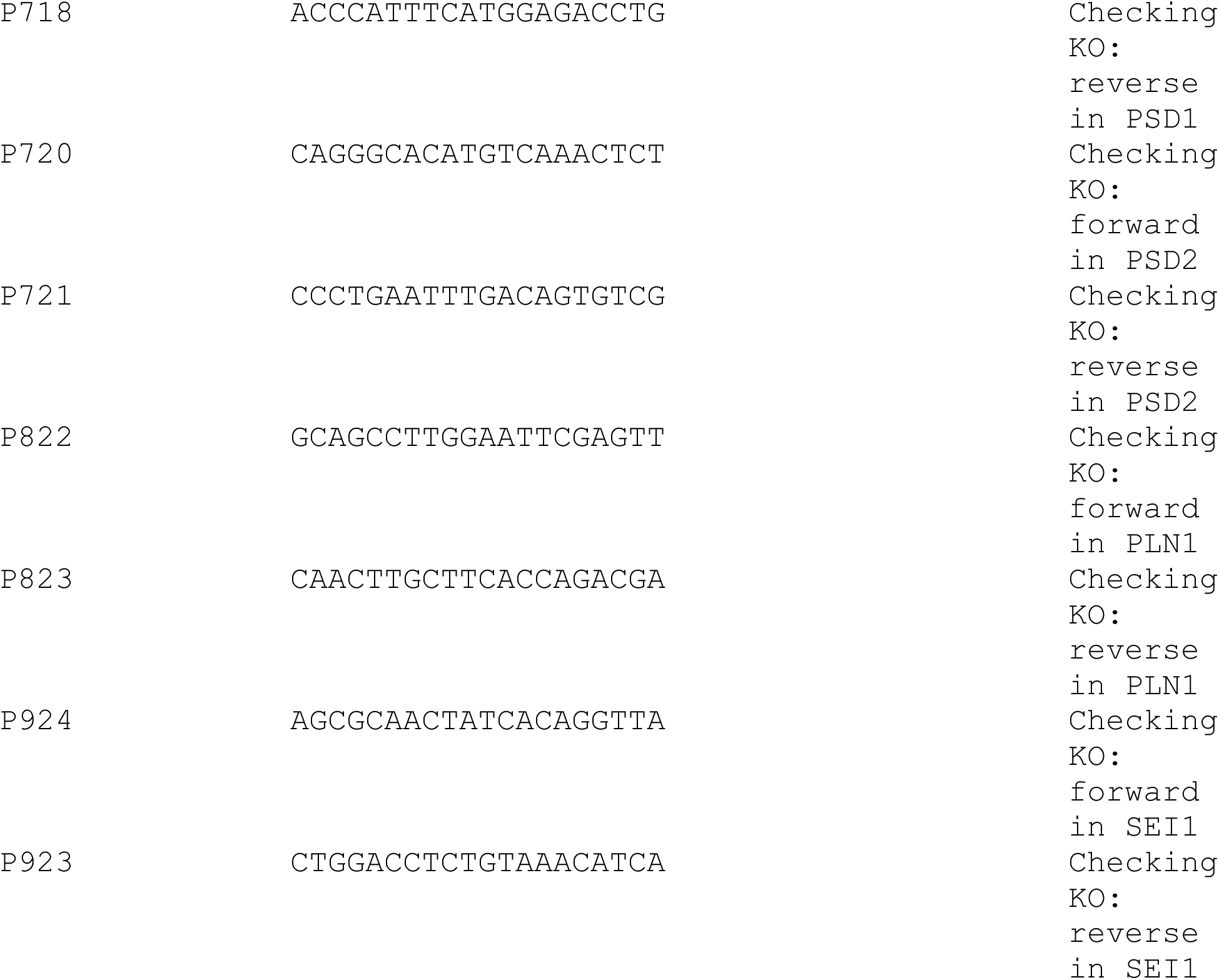
Primer List:

**Table S2.** Strain list:

| Strains | Genotype | Source |
| --- | --- | --- |
| ATY1842 | W303 *lro1::URA3 are1::HIS3 are2::LEU2*<br>*TRP1-GAL1-10 (pr) -DGA1 pAGL-ClONAT* | J. Friedman |
| DBY116 | W303 *lro1::URA3 are1::HIS3 are2::LEU2*<br>*TRP1-GAL1-10 (pr) -DGA1 pAGL-ClONAT*<br>*psd1::KanMX4 psd2::HyG* | This study |
| DBY151 | W303 *lro1::URA3 are1::HIS3 are2::LEU2*<br>*TRP1-GAL1-10 (pr) -DGA1 pAGL-ClONAT*<br>*pln1::KanMx4 sei1::HygMX* | This study |

**Figure S1.**
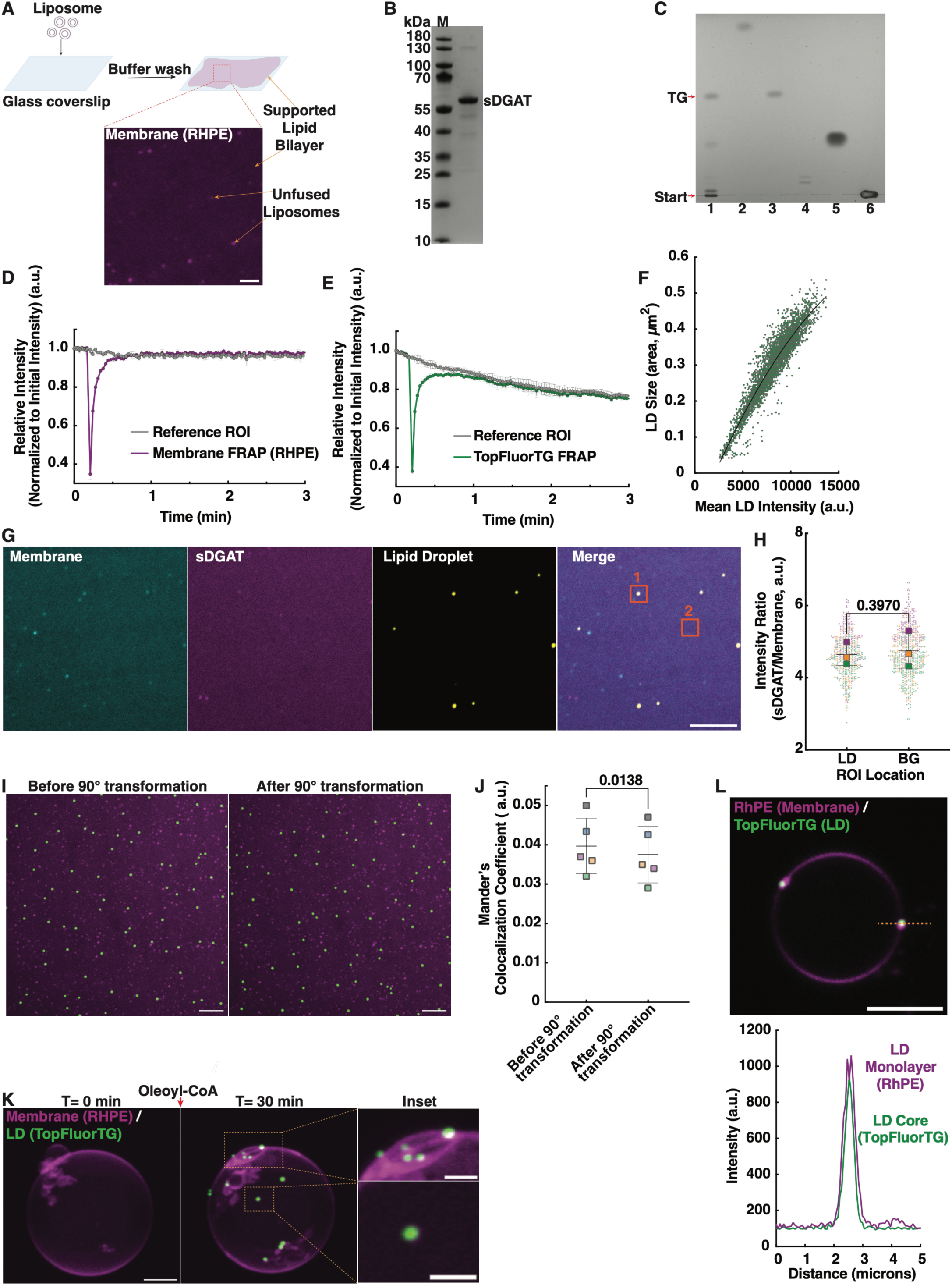
**Supplemental data for Fig. 1** A. Cartoon of SLB formation (top). View of RHPE-containing SLB (bottom). B. SDS-PAGE analysis of purified sDGAT and stained with Coomassie. M = molecular weight markers. C. Lipids extracted from liposomes containing 10% DG and incubated with sDGAT and oleoyl-CoA for 2 h at 30°C (lane 1) were separated by thin layer chromatography (TLC). Lipid standards in other lanes: wax ester (lane 2), TG (lane 3), DG (lane 4), oleic acid (lane 5), and DOPC (lane 6). The TLC plate was developed with hexane:diethyl ether:acetate (80:20:2) and lipids visualized by staining with copper sulphate solution. D. and E. FRAP of fluorescent lipids on LDs made in SLBs with DOPC:DG:DOGS-NiNTA:RHPE:TopFluorTG (87.3:10:2.5:0.1:0.1 mol%). Mean +/− std. dev. from 3 independent experiments. F. Graph of mean intensity and size of ∼3800 LDs formed on SLBs with DOPC:DG:DOGS-NiNTA:TopFluorTG (87.4:10:2.5:0.1 mol%) after incubation with oleoyl-CoA at 30°C for 1 hr. G. Image of SLBs with DOPC:DG:DOGS-NiNTA (87.4:10:2.5 mol%) and trace amounts (0.1 mol%) of 18:1 TopFluor-PE (yellow) after incubation with oleoyl-CoA for at 30°C for 1 h. The SLBs contained 1000nM sDGAT-mCherry (magenta) and were stained with Lipi-Blue (Dojindo; cyan) to visualize LDs. Merge shows representative LD (box 1) and background (box 2) used for quantitation in G. Bar = 2µm. H. Quantification of experiments as in panel F. Intensity of sDGAT-mCherry signal of regions of interest (ROIs) with LDs and ROIs without LDs, background (BG), were determined. Colored squares show the ratio of the mean fluorescence signal of ROIs with an LD to a BG ROI calculated from at least 150 LD ROIs per independent experiment; dots with the same color as the squares were used to calculate the mean. Bars and lines show the mean and std. dev. of the three experiments; p value calculated from Welch’s t-test. I. J. SLBs with DOPC:DG:DOGS-NiNTA:TopFluorTG (87.4:10:2.5:0.1 mol%) were imaged before (magenta) and after (green) incubation with oleoyl-CoA at 30°C for 5 min. Contrast of the images was adjusted so that unfused liposomes on top of the SLBs were visible. (J) Manders’ coefficients were calculated from co-localization of LDs and unfused liposomes before and after 90° rotation of the green image. Mean +/− std. dev. for 5 independent experiments, p value calculated from paired t-test. K. Images from a movie (Movie 3) showing LD formation on GUVs with DOPC:DG:DOGS-NiNTA (93.2:4:2.5 mol%) and 0.1 mol% of TopFluorTG (green) and 0.2 mol% RHPE (magenta). Bar = 5 μm, inset = 2 μm. L. Fluorescence signal along dotted line of signal from LD formed as in Fig. 1G. Bar = 5 μm.

**Figure S2.**
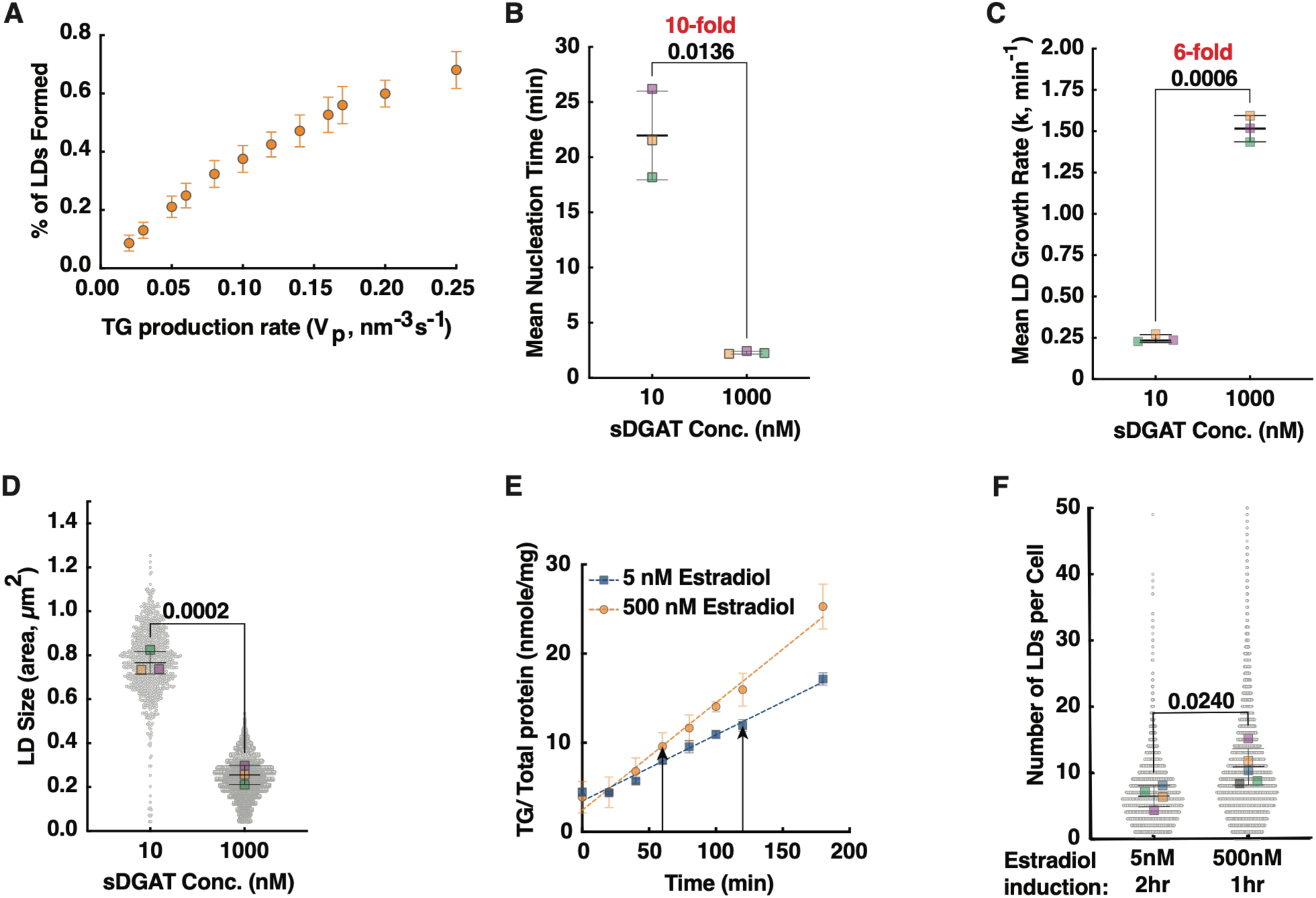
**Supplemental data for Fig. 3** A. Number of LDs formed in Monte Carlo-like simulations performed with different velocities of TG addition (V_p_) and constant line and surface tension. % of LDs formed is the number of possible LD nucleation sites that contain LDs at the end of the simulations. Mean and std. dev. of 3 independent experiments. B. C. Mean nucleation time (B) and growth rate (C) of LDs formed in assays as in Fig. 3C. Nucleation time is the first frame in which a new LD is detected. Squares are means of independent experiments for each condition; at least 10 LDs (10 nm sDGAT) and 290 LDs (1000 nm sDGAT). Bars are the mean and std. dev. of 3 independent experiments; p value calculated from Welch’s t-test. D. Size of LDs formed 1 h after oleoyl-CoA addition in assays as in Fig. 3C. Gray dots are individual LDs, colored squares are means of independent experiments with at least 900 LDs for 10 nM sDGAT and 16,000 LDs for 1000 nM sDGAT. Bars are the mean and std. dev. of 3 independent experiments. p value calculated from Welch’s t-test. E. Quantification of TG in *GAL1-DGA1* 3Δ cells after estradiol addition to the to medium. Mean and std. dev. of three independent experiments. Arrows indicate TG level at 1h for 500 nM estradiol and 2 h for 5 nM estradiol. F. Number of LDs formed in the yeast strain *GAL1-DGA1* 3Δ after *DGA1* expression was induced with the indicated amount of estradiol. Gray dots indicate number of droplets in individual cells; colored squares are means of independent experiments with at least 1500 cells per condition. Bars and lines show the mean and std. dev. of the three experiments; p value calculated from Welch’s t-test.

**Figure S3.**
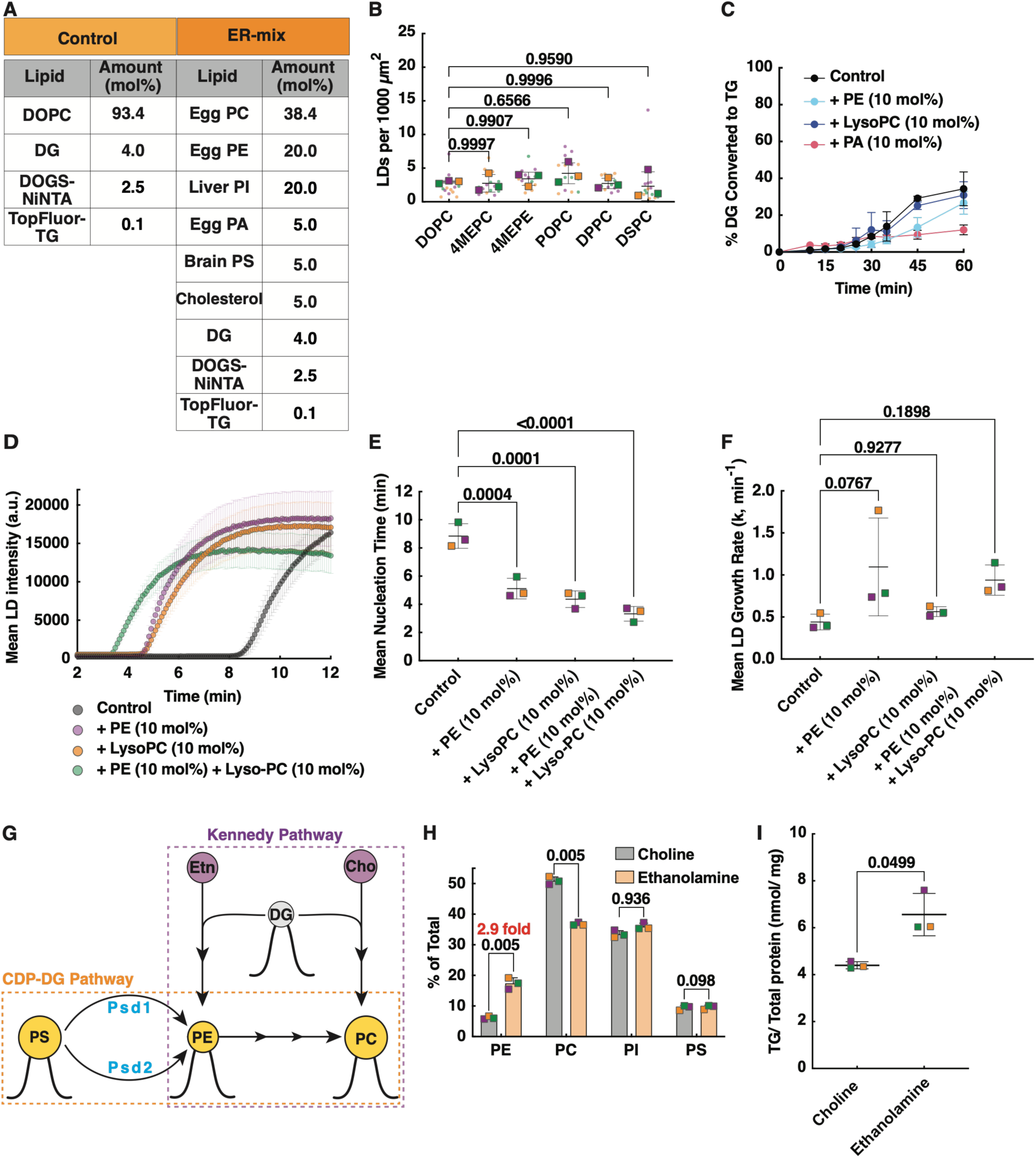
**Supplemental data for Fig. 3**. A. Lipid Composition for ER-like and control lipid mixes. B. LD density for LD formation experiment with SLBs using the control lipid mix (panel A) and 10 mol% of the indicated lipids. The results are depicted as described in Fig. 3F. C. Liposomes were prepared with DOPC (control), the indicated additional lipid, and 4 mol% [^3^H]DG. The enzyme sDGAT (1000 nM) was added and reactions were started with oleoyl-CoA addition (time = 0). The percent of [^3^H]DG converted to TG was determined. D. Mean intensity over time of LDs formation on SLBs DOPC, 4 mol% DG, 10 mol% of the indicated lipids, and 1000 nM sDGAT. Oleoyl-CoA was added at time = 0. Mean +/− std. dev. of 780 LDs. E. F. Mean nucleation time (E) and growth rate (F) of LDs formed in assays as in panel D. Nucleation time is the first frame in which a new LD is detected. Squares are means of independent experiments; at least 100 LDs were analyzed per condition. Bars are the mean and std. dev. of 3 independent experiments; p values calculated from one-way ANOVA. G. Cartoon depicting the two biosynthesis pathways in yeast that produce PE and PC. H. The amount of TG in the cells shown in Fig. 3G. Mean and std dev. of 3 independent experiments. p value calculated from Welch’s t-test. I. GAL1-DGA1 3Δ *psd1*Δ *psd2*Δ cells were labeled to steady state with [^3^H]palmitate while grown as in Fig. 3H. Lipids were extracted and quantified. Mean and std dev. of 3 independent experiments; p values calculated from paired t-tests.

**Figure S4.**
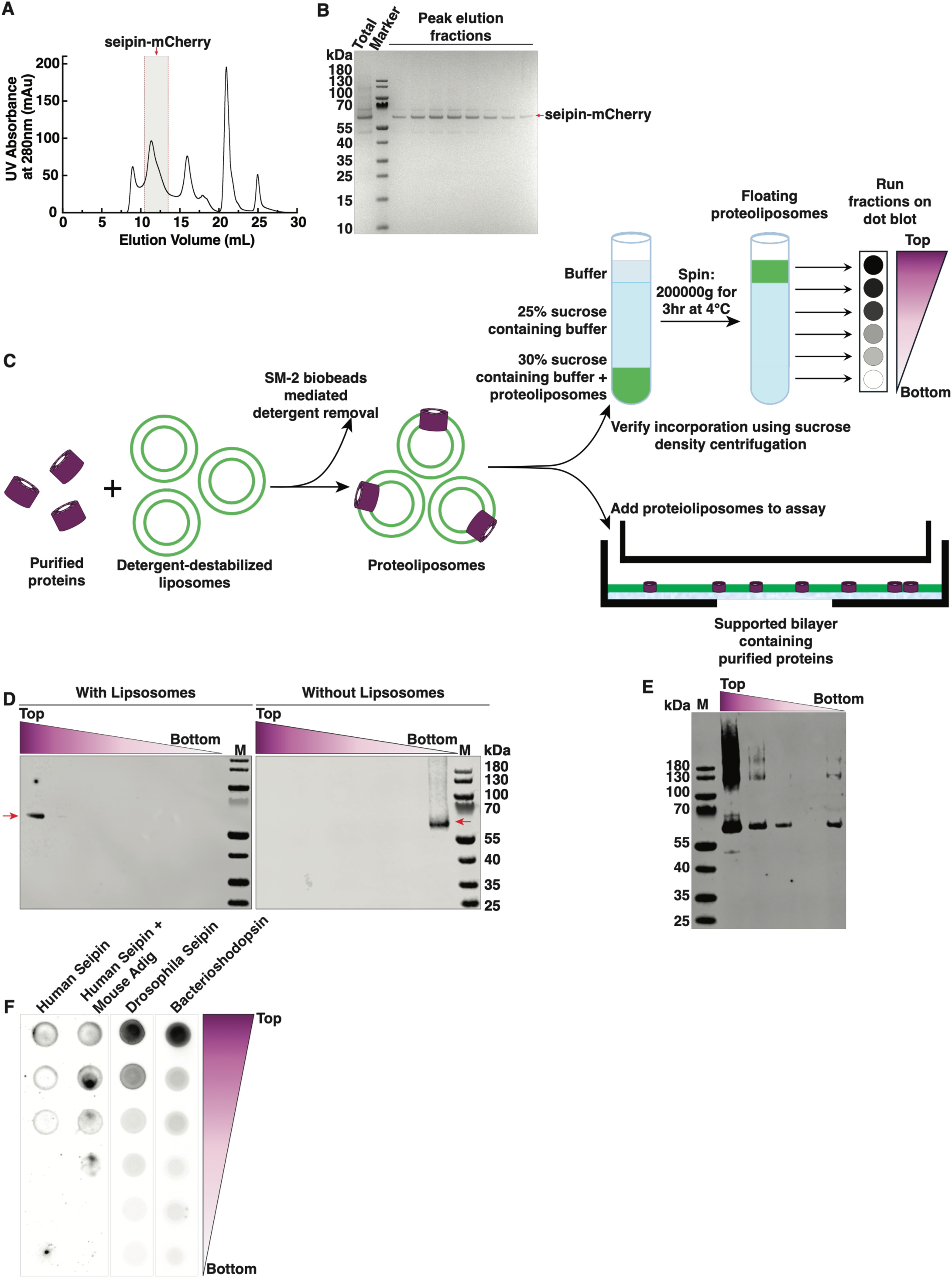
**Supplemental data for Fig. 4**. A. Size exclusion chromatography (SEC) profile of *Drosophila* seipin-mCherry oligomers. Gray-shaded area indicates peak elution fractions containing seipin oligomers. B. SDS-PAGE analysis of seipin preparation shown in panel A. Protein in the pooled sample before SEC (Total) and peak elusion fractions after SEC as they eluted from the column (gray shaded area in panel A) are shown. Gel was stained with Coomassie. M = molecular weight markers. Red arrow indicates location of purified seipin–mCherry. C. Scheme for incorporation of purified proteins into proteoliposomes (*see Methods for details*). Briefly, purified proteins were mixed with detergent-destabilized liposomes and incubated with SM-2 biobeads for gradual detergent removal and proteoliposomes formation. Protein incorporation was verified using sucrose density centrifugation (top). Proteoliposomes were used to form SLBs containing purified proteins (bottom). D. Western blot of *Drosophila* seipin–mCherry proteoliposomes (left) or on controls lacking liposomes (right) fractionated using sucrose density gradient centrifugation. Red arrow indicates the protein. E. Western blot of *Drosophila* seipin-mCherry proteoliposomes fractions after carbonate extraction and sucrose density gradient centrifugation. F. Dot blot of proteoliposomes made with the indicated proteins after fractionation using sucrose density gradient centrifugation.

**Figure S5.**
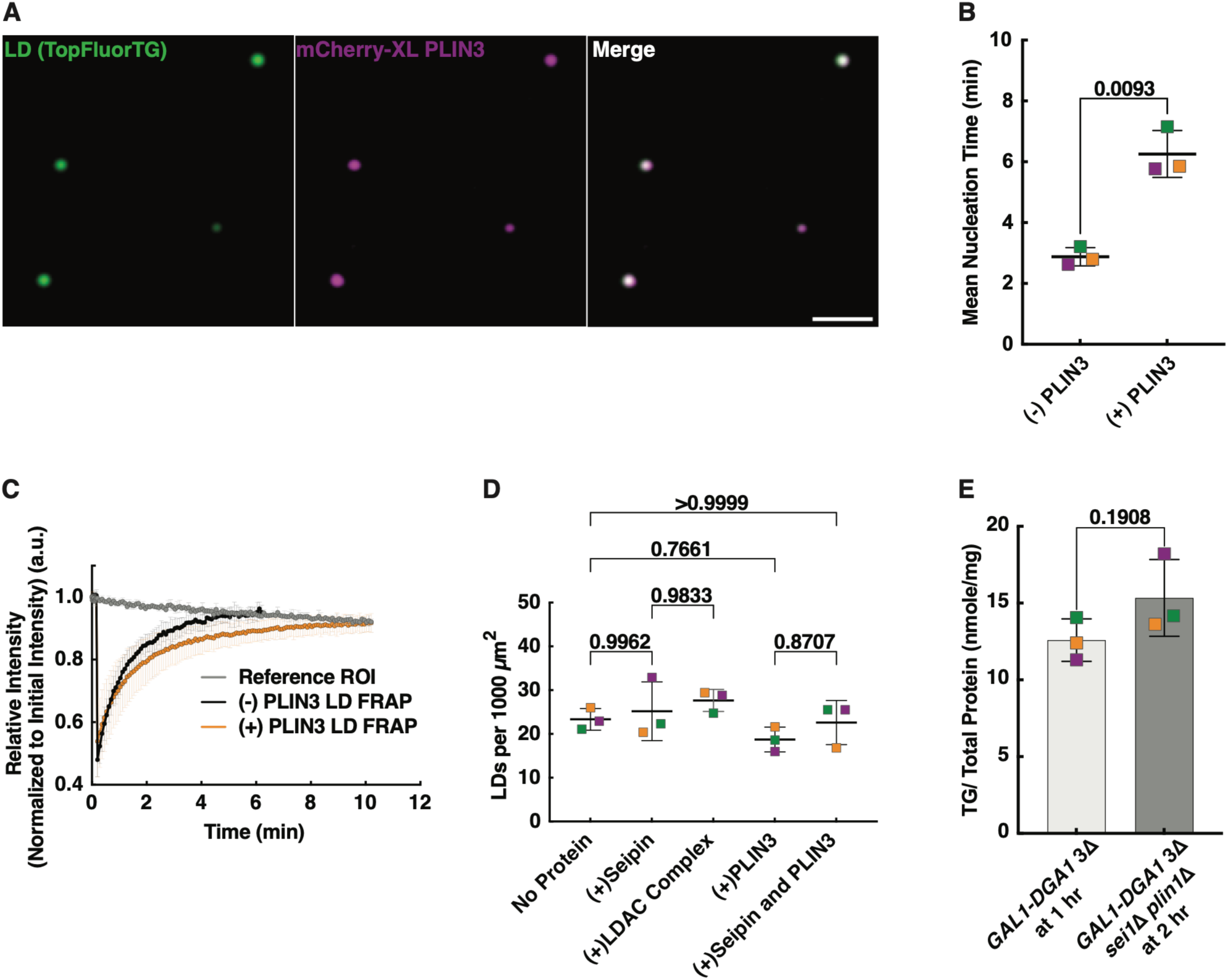
**Supplemental data for Fig. 5**. A. Representative image of mCherry-XL-PLIN3 localized to LDs. Bar = 2µm. B. Mean nucleation time for LD formation with and without PLIN3. Squares show the mean of at least 100 LDs. Bars are the mean and std. dev. of 3 independent experiments; p values calculated from Welch’s t-tests. C. FRAP of TopFluorTG signal in LDs formed with or without 250 nM mCherry-XL PLIN3). Mean +/− st. dev. of 3 independent experiments. D. The amount of TG in the cells shown in Fig. 5E. Mean and std dev. of 3 independent experiments; p value calculated from Welch’s t-test. E. Amount of TG produced in the yeast strains shown in Fig. 5I grown in the same conditions.

## Extended Data

**Movies 1-4**

### Legends for movies in Extended Data

**Movie 1.** Droplets forming on an a SLB with DOPC:DG:DOGS-NiNTA:TopFluorTG (87.4:10:2.5:0.1 mol%) and 1000 nM sDGAT. Oleoyl-CoA added at start of movie. Images taken every 5 seconds for 5 minutes. Bar = 5μm

**Movie 2.** Droplets forming on a GUV with DOPC:DG:DOGS-NiNTA:TopFluorTG:RHPE (93.3:4:2.5:0.1:0.2 mol%). A z-stack of 21 images with step size of 0.3 µm was taken every 5 minutes for 30 minutes. Oleoyl-CoA added at start of movie. Bar = 5µm

**Movie 3.** Droplets forming in Cos-7 cells. 400µM oleic acid added at the start of the movie. Images taken every 20 seconds for 35 minutes. Bar = 10µm

**Movie 4**. An SLB containing DOPC:DG:DOGS-NiNTA:TopFluorTG (87.4:10:2.5:0.1 mol%) and 1000 nM sDGAT was incubated for ∼1h at 30°C with 1.8 µM oleoyl-CoA. A second wave of LD formation was initiated at time = 0 by adding 18 µM oleoyl-CoA. Bar = 2µm.

### Codon optimized seq of ALFA-tagged *Drosophila* LDAF1

CCGAGCCGTCTGGAAGAAGAACTGCGCCGTCGCCTGACCGAACCGAGCAGCGGCCTGGTGCCGCGCGGCAGCATGGCGCATTCCAACAAAGCGCGCAAGGCGGCTAAAGAAGCAAACGCAAAGAAATCGGCCGGTTCTGGCACTGAATCAAAGCCTGAACATGTTAAAATCCATGGCCCCGGTGACCGCTCAGCGGGCGTGCTGACCGGAGATCAATTCTTAGAAAGCCTGTTACAGACTCTGTATCGCTCGTGGTTACAAATCTGTCGTCTTGCCGATTTCGTAATGAGCGAACTTGGGGGCAATGATTTTTTGGAGGCGACCGCGGCGTGGTGCACACGCAATCCACATGTTGCGATTTGCTTGCTGGCCGGGGGCCTGGTGTTTATGCTGCCTTTTCTGATCATTTTCGGCTTCGGCATTGCCACTATGGTTATGACGTTCACCGGCTTGTTGGTGCTGGAAGGAACTCTGCTGACCATTGTAAGTATGGTTTTTTTCGCCTGCCTTGGAGGTCTGGTGATCATGGTTCCATTATTCGGCGTTGCCGCCGTAGCCGCGTACTTCGGCTTCGCACAAGTTTACGGCCTGTGCGACGGCATTGAAATCCACAAATCCGCCTGGGTGATGTTCTTTCGCGACCAGCGTAAAGCGGGCGATCCTCCACCGGCGGCGGTGACCATTACTGACGAACCGACTAGCACTACGGCG

